# Immune-responsive biodegradable scaffolds for enhancing neutrophil regeneration

**DOI:** 10.1101/2022.01.21.477275

**Authors:** Matthew D. Kerr, David A. McBride, Wade T. Johnson, Arun K. Chumber, Alexander J. Najibi, Bo Ri Seo, Alexander G. Stafford, David T. Scadden, David J. Mooney, Nisarg J. Shah

**Affiliations:** Department of Nanoengineering, University of California, San Diego, La Jolla, CA 92093, USA; Chemical Engineering Program, University of California, San Diego, La Jolla, CA 92093, USA; John A. Paulson School of Engineering and Applied Sciences, Harvard University, Cambridge MA 02138; Wyss Institute for Biologically Inspired Engineering, Harvard University, Cambridge, Massachusetts 02138; Department of Stem Cell and Regenerative Biology, Harvard University, Cambridge, MA 02138, USA; Harvard Stem Cell Institute, Cambridge, MA 02138, USA; Center for Regenerative Medicine, Massachusetts General Hospital, Boston, MA 02114, USA; Program in Immunology, University of California, San Diego, La Jolla, CA 92093, USA

## Abstract

Neutrophils are essential effector cells for mediating rapid host defense and their insufficiency arising from therapy-induced side-effects, termed neutropenia, can lead to immunodeficiency-associated complications. In autologous hematopoietic stem cell transplantation (HSCT), neutropenia is a complication that limits therapeutic efficacy. Here, we report the development and in vivo evaluation of an injectable, biodegradable hyaluronic acid (HA)-based scaffold, termed HA cryogel, with myeloid responsive degradation behavior. In mouse models of immune deficiency, we show that the infiltration of functional myeloid-lineage cells, specifically neutrophils, is essential to mediate HA cryogel degradation. Post-HSCT neutropenia in recipient mice delayed degradation of HA cryogels by up to 3 weeks. We harnessed the neutrophil-responsive degradation to sustain the release of granulocyte colony stimulating factor (G-CSF) from HA cryogels. Sustained release of G-CSF from HA cryogels enhanced post-HSCT neutrophil recovery, comparable to pegylated G-CSF, which, in turn, accelerated cryogel degradation. HA cryogels are a potential approach for enhancing neutrophils and concurrently assessing immune recovery in neutropenic hosts.

## Introduction

Neutrophils mediate essential host defense against pathogens and are among the earliest responders in tissue injury ^1–3^. Neutrophil deficiency, termed neutropenia, contributes to opportunistic infections and could impair tissue regeneration in affected individuals ^4–7^. In autologous hematopoietic stem cell transplantation (HSCT) pre-conditioning myelosuppressive regimens can contribute to a marked transient post-therapy impairment of neutrophils and render recipients susceptible to immune deficiency-associated complications for up to several weeks ^6, 8–11^.

Post-HSCT neutrophil regeneration follows successful bone marrow engraftment of transplanted hematopoietic cells ^12, 13^, facilitated by granulocyte colony stimulating factor (G-CSF)-mediated granulopoiesis of hematopoietic cells ^14–16^. Neutropenia is typically treated as an emergency and, in a subset of patients, the risk of neutropenia may be prophylactically addressed with post-HSCT subcutaneous injection of recombinant human G-CSF (filgrastim) to facilitate recovery ^6, 14, 17, 18^. Daily injections are used as G-CSF has a half-life of a 3 – 4 hours, which can be extended by conjugating G-CSF with polyethylene glycol (PEGylation) ^19, 20^. However, immune responses against PEG have been demonstrated to enhance clearance of PEG-G-CSF in an antibody-dependent manner ^21^. As multiple cycles of PEG-G-CSF treatment are common, long-term treatment could be rendered ineffective. Therefore, the development of a sustained release method to deliver G-CSF while avoiding immune responses against PEG, and concurrently assess neutrophil function could greatly improve the current standard-of-care.

Seeking to improve post-HSCT recovery of neutrophils and simultaneously assess recovery, we developed a biodegradable depot to prophylactically deliver G-CSF in post-HSCT recipients. The depot comprised a porous injectable scaffold made by low-temperature crosslinking, termed cryogelation, of hyaluronic acid (HA), an easily sourced and readily derivatized anionic glycosaminoglycan, termed ‘HA cryogel.’ As a component of the extracellular matrix, endogenous HA is a substrate for degradation by myeloid cells through enzymatic action and by neutrophil-mediated oxidation ^22–24^. Harnessing the immune-responsiveness of HA, we characterized in vivo degradation of HA cryogels in immune deficient and post-HSCT mice and identified myeloid cell infiltration in HA cryogels to be key mediators in facilitating degradation, which was significantly reduced or altogether eliminated in mice with severely deficient neutrophil function. Transient but profound post-HSCT myeloid depletion significantly delayed degradation of HA cryogels until recovery of neutrophils ^25^. As the degradation profile of HA cryogels was responsive to neutrophil recovery, we harnessed encapsulated G-CSF to facilitate the sustained release, which was mediated by HA cryogel degradation. Neutrophil reconstitution was enhanced in post-HSCT mice injected with G-CSF-encapsulated HA cryogels, comparable to a single dose of PEGylated G-CSF, which accelerated HA cryogel degradation.

## Results

### Synthesis and characterization of HA cryogels

Click-functionalized HA was prepared by conjugating either tetrazine (Tz) amine or norbornene (Nb) methylamine to HA using carbodiimide chemistry. Nb-functionalized HA (HA-Nb) was reacted with Tz-Cy5 to form Cy5-labeled HA-Nb (Cy5-HA-Nb) (**Fig. 1a**). Tz amine-functionalized HA (HA-Tz) was prepared at 7% degree of substitution (termed high-DOS). 0.8% DOS HA-Tz (termed low DOS) was also prepared for comparison. Endotoxin levels of HA-Tz and Cy5-HA-Nb were quantified to be less than 5 endotoxin units/kg, the threshold pyrogenic dose for preclinical species (**Supplementary Table 1**) ^26^. To maximize polymer concentration while maintaining proper viscosity to achieve mixing, 0.6% w/v aqueous solutions of HA-Tz and Cy5-HA-Nb, pre-cooled to 4°C, were well mixed in a 1:1 (v/v) ratio by vortexing (**Fig. 1a**). The solution was then pipetted onto individual pre-cooled (-20°C) cryomolds (30 μL/mold) and immediately transferred to a -20°C freezer and allowed to freeze (**Supplementary Note 1**), to generate Cy5-HA cryogels (**Fig. 1b, c**).

**Fig. 1.**
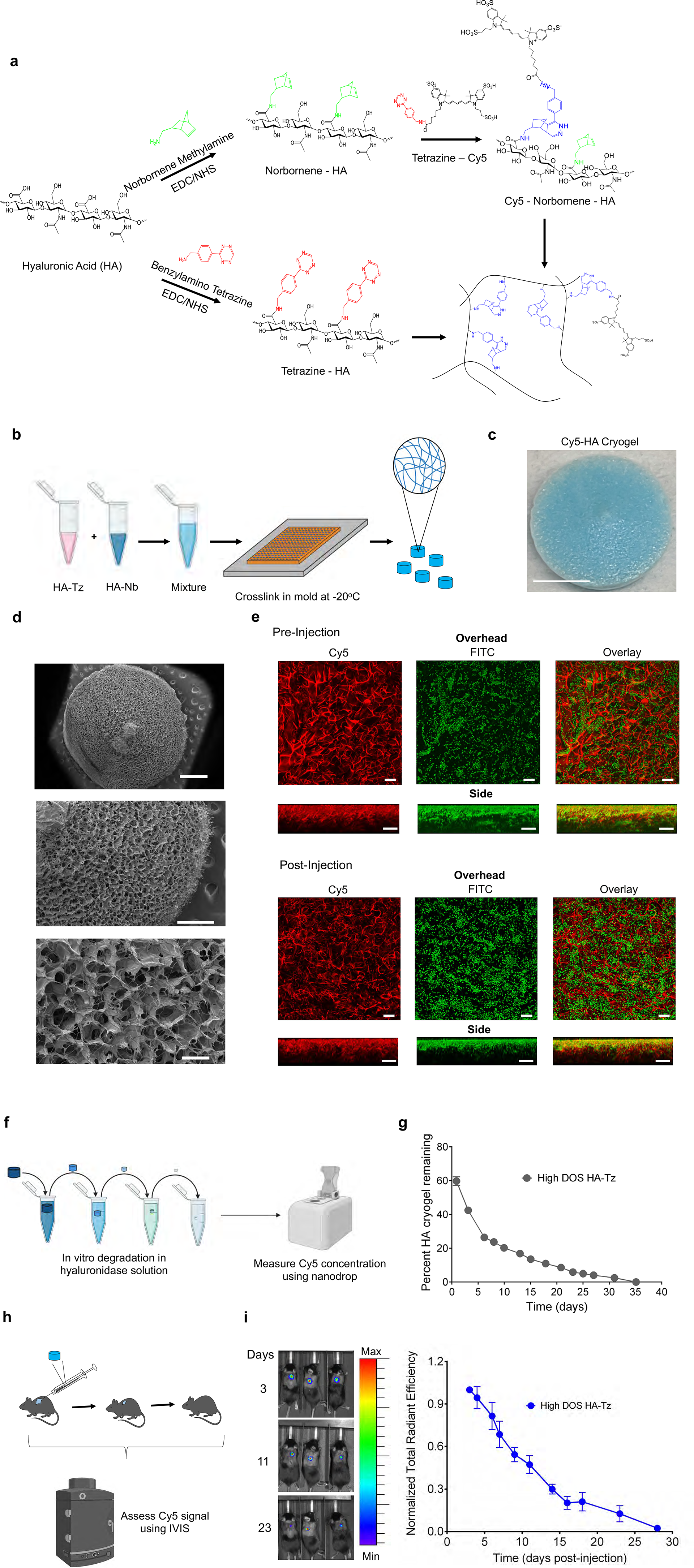
Production and characterization of Cy5-HA cryogel (**a**) Schematic for tetrazine (Tz) and norbornene (Nb) functionalization of HA, Cy5 functionalization of Nb functionalized HA (Cy5-HA-Nb) and crosslinking of Tz functionalized HA with Cy5-HA-Nb. (**b**) Schematic for producing Cy5-HA cryogels. (**c**) Representative photograph of lyophilized Cy5-HA cryogel. Scale bar = 1mm. (**d**) Representative SEM image depicting Cy5-HA cryogels. Top scale bar = 1mm, middle scale bar = 500μm, bottom scale bar = 100μm. (**e**) Confocal microscopy image, overhead and side views, depicting hydrated Cy5-HA cryogels pre-and post-injection incubated with 10μm FITC-labeled microparticles. Scale bar = 100μm. (**f**) Schematic depicting workflow for in vitro Cy5-HA cryogel degradation study. (**g**) Measuring Cy5-HA cryogel degradation in vitro by quantifying the Cy5-signal in supernatant at pre-determined timepoints normalized to total Cy5-signal in supernatant across all timepoints. (**h**) Schematic depicting workflow for in vivo Cy5-HA cryogel degradation study. (**i**) Representative in vivo imaging system (IVIS) fluorescence images of gel degradation in mice and measuring Cy5-HA cryogel degradation in vivo by quantification of total radiant efficiency normalized to initial day 3 timepoint. IVIS Images are on the same scale and analyzed using Living Image Software. Data in **g** represents mean ± s.d. of n=4 HA cryogels. Data in **i** represents mean ± s.e.m. of n=4 HA cryogels. Part of figure **1b, f and h** were created with BioRender.com.

To characterize Cy5-HA cryogels, we estimated the swelling ratio by comparing the hydrated vs. cast volume and the aqueous mass composition from the wicked mass and fully hydrated mass. The swelling ratio was 1.5 ± 0.1 in both low- and high-DOS Cy5-HA cryogels (**Supplementary Fig. 1a**). The aqueous mass composition was 76.3 ± 4.0% and 71.9 ± 2.6% in low- and high-DOS Cy5-HA cryogels respectively (**Supplementary Fig. 1b**).

To measure surface porosity of lyophilized Cy5-HA cryogels, we used scanning electron microscopy (SEM) (**Fig. 1d, Supplementary Fig. 1c**). The surface pore structure images were used to measure the average pore diameter using FIJI, which were between 80-180μm and 40-90μm for low- and high-DOS Cy5-HA cryogels respectively (**Supplementary Fig. 1d**). To characterize interconnectedness of the Cy5-HA cryogel pore structure, we incubated fully hydrated low- and high-DOS Cy5-HA cryogels with Fluorescein isothiocyanate (FITC)-labeled 10μm diameter melamine resin particles and imaged using a confocal microscope (**Fig. 1e, Supplementary Fig. 1e**). Since the route of administration of the Cy5-HA cryogels is through a needle, we repeated this experiment with Cy5-HA cryogels after injection and observed similar penetration of the FITC-labeled 10μm particles (**Fig. 1e, Supplementary Fig. 1e**). Image analysis of z-stacked images showed co-localization of the FITC-labeled 10μm particles with Cy5-HA up to a depth of 100μm below the surface, which was the limit of detection (**Supplementary Fig. 1f**). Both low- and high-DOS Cy5-HA cryogels maintained pore morphology and relative surface pore size distribution following lyophilization and rehydration (**Supplementary Fig. 1g, 1h**). Cy5-HA cryogels also maintained shape and structure post-injection (**Supplementary Movie 1**).

To confirm susceptibility of Cy5-HA cryogels to enzymatic degradation, we used a hyaluronidase-2 (HYAL2)-based in vitro assay (**Fig. 1f**). In native HA, HYAL2 cleaves internal beta-N-acetyl-D-glucosaminidic linkages resulting in fragmentation of HA ^27^. Here, HYAL2 degraded HA cryogels and high DOS Cy5-HA cryogels degraded at a slower rate compared to the low DOS Cy5-HA cryogels in vitro (**Fig. 1g, Supplementary Fig. 2a**). To confirm in vivo degradation, low- and high-DOS Cy5-HA cryogels were injected in subcutaneously in the hind flank of C57Bl/6J (B6) mice and degradation was measured using in vivo imaging system (IVIS) fluorescence spectroscopy (**Fig. 1h**). In contrast to in vitro degradation, both low- and high-DOS Cy5-HA cryogels degraded at a similar rate (**Fig. 1i, Supplementary Fig. 2b**). This observation, together with the finding of a similar pore size distribution in hydrated low- and high-DOS HA cryogels (**Supplementary Fig. S1h**), supported the selection of one of the types of HA cryogels for subsequent experiments, and we selected high-DOS HA cryogels. To characterize if HA cryogels made from different batches of derivatized HA affected in vivo degradation, we compared degradation of Cy5-HA cryogels made from three distinct batches of Cy5-HA-Nb and HA-Tz and confirmed that all Cy5-HA-cryogels degraded at a similar rate (**Supplementary Fig. 2c**).

### Depletion of immune cell subsets affects cellular infiltration into HA cryogels

As the HSCT pre-conditioning regimen depletes all immune cell lineages, we first sought to measure the effect of immune depletion on HA cryogel degradation. Cy5-HA cryogels were subcutaneously injected into the hind flank of untreated B6 mice (**Fig. 2a**) and the degradation profile was compared to that in B6 mice receiving (i) anti-Ly6G antibodies and anti-rat κ immunoglobulin light chain to deplete neutrophils (**Fig. 2b**), (ii) clodronate liposomes to deplete macrophages (**Fig. 2c**), (iii) anti-CD4 and anti-CD8 antibodies to deplete T cells (**Supplementary Fig. 3a**), (iv) anti-B220 to deplete B-cells (**Supplementary Fig. 3b**) and immune deficient NOD.Cg-*Prkdc^scid^ Il2rg^tm1Wjl^*/SzJ (NSG) mice (**Fig. 2d**). The durability of depletion was assessed by measuring peripheral blood cellularity throughout the duration of the degradation study (**Supplementary Fig. 3c-l, Supplementary Table 2**). In untreated immune competent mice, the average half-life of Cy5-HA cryogels, quantified as the time to achieve a 50% reduction in fluorescence intensity, was about 9.5 days (**Supplementary Fig. 3m**). The average half-life in the macrophage, neutrophil, T cell, and B cell depleted mice was similar at about 11.8 days, 11.3 days, 9.6, and 10.2 respectively (**Supplementary Fig. 3m**). In contrast, only a 35% reduction in Cy5 signal intensity was measured after 3 months in the NSG mice (**Fig. 2d**). Retrieval of Cy5-HA cryogels from sacrificed mice at the endpoint confirmed that the gels had minimally degraded (**Supplementary Fig. 3n**).

**Fig. 2.**
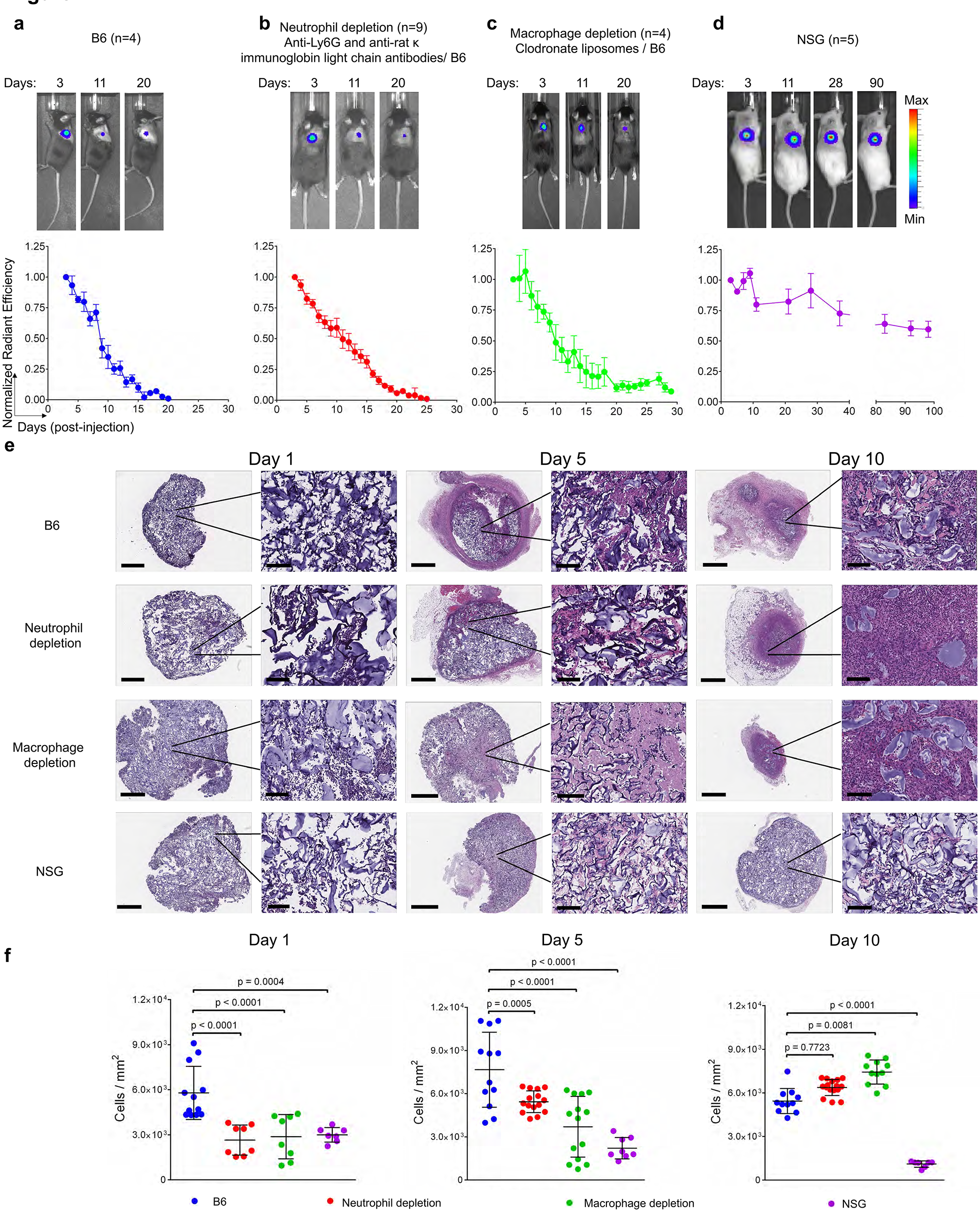
Cy5-HA cryogel degradation in immunodeficient mice Representative IVIS fluorescence images of Cy5-HA cryogel degradation and quantification by measuring total radiant efficiency normalized to initial day 3 timepoint of (**a**) untreated B6 mice, (**b**) neutrophil depleted B6 mice, (**c**) macrophage depleted B6 mice, and (**d**) NSG mice. IVIS Images are on the same scale and analyzed using Living Image Software. (**e**) Hematoxylin and eosin (H&E) stained histological sections of explanted Cy5-HA cryogels from the above groups, at days 1, 5, and 10 post-injection. Full view scale bar = 800μm, magnified scale bar = 100μm. (**f**) Quantification of cellular density in the sections from **e**. Data in **a**-**d** represent mean ± s.e.m. of n=4-9 and are representative of at least two separate experiments. Data in **f** represents mean ± s.d. of n=7-12 and were compared using student’s t-test.

To assess cellular infiltration and the foreign body response, Cy5-HA cryogels were explanted from the above groups at 1-, 5-, and 10-days post-injection and stained using haemotoxylin and eosin (H&E). In Cy5-HA cryogels retrieved from all groups except NSG mice, the total cellularity increased from day 1 to 10 and formed a distinct capsule encapsulating the HA cryogel, indicative of a foreign body response (**Fig. 2e, Supplementary Fig. 4a**). In NSG mice, some infiltrates were quantified on day 1, however there was no appreciable increase in cellularity at the later timepoints or a capsule by day 10 (**Fig. 2e**). H&E slides were further analyzed to quantify the cell density in the different groups. Differences in infiltrates between the untreated and all immunodepleted B6 mice were significant at the earlier timepoints, and either increased or remained constant in all immunodepleted B6 (**Fig 2f, Supplementary Fig. 4b**). In contrast, cell infiltrates in HA cryogels retrieved from NSG mice reduced steadily and were 80% lower than untreated B6 mice by day 10 (**Fig. 2f**).

To identify the immune cells contributing to HA cryogel degradation, we quantified cell infiltrates in the Cy5-HA cryogels 1- and 10-days post-injection using flow cytometry in untreated and immune depleted B6 and NSG mice (**Fig. 3a, b, Supplementary Fig. 5a, b**). Viability of infiltrating cells, quantified by negative AnnexinV staining, was consistently greater than 95% in all groups (**Supplementary Fig. 5c**).

**Fig. 3.**
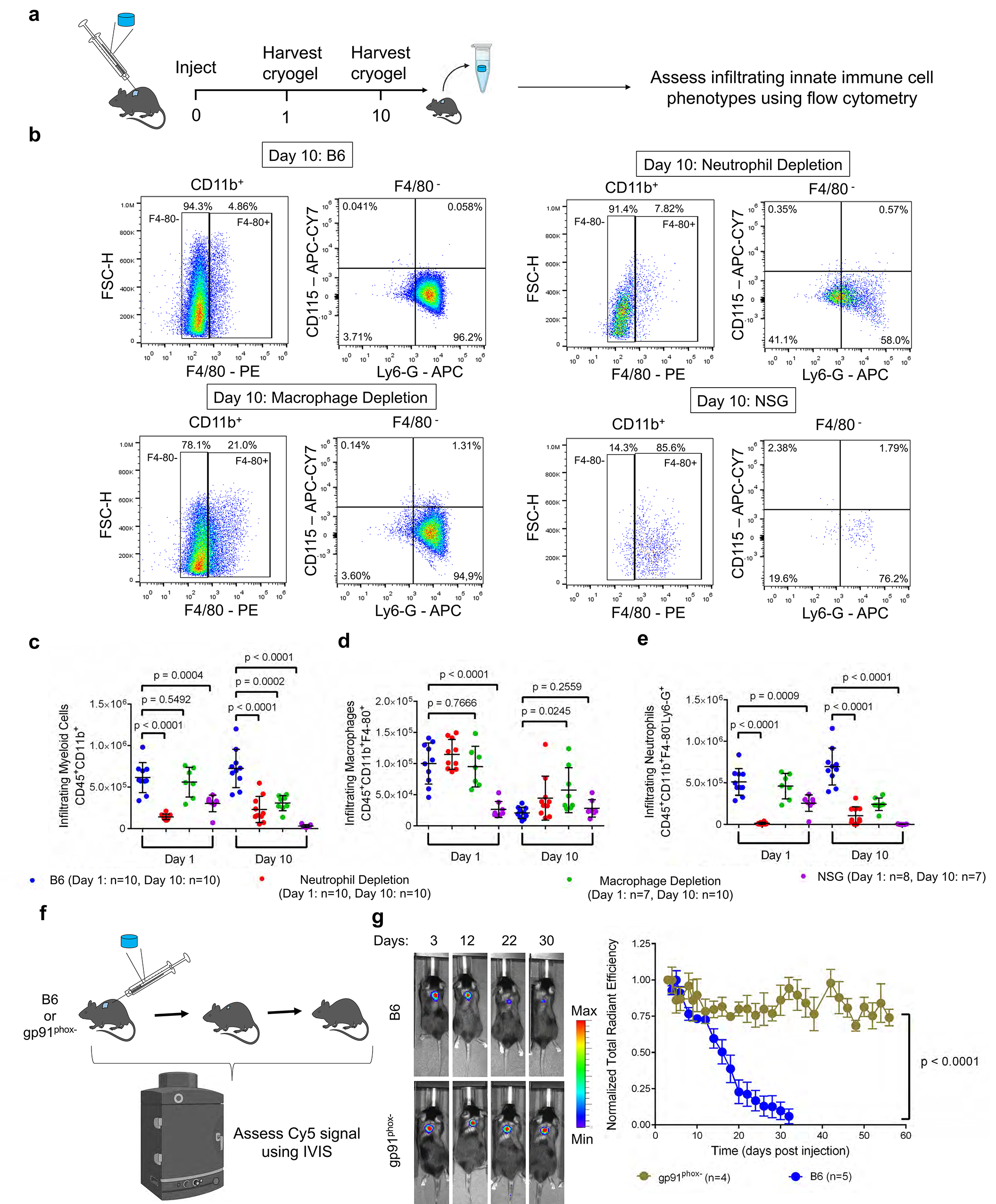
Assessment of innate immune cell infiltration into Cy5-HA cryogels (**a**) Schematic for the quantification of innate immune cell content in Cy5-HA cryogels. (**b**) Representative flow cytometry plots depicting gating strategy to determine cellular identity of CD45^+^ CD11b^+^ F4/80^+^ (macrophage) cells, CD45^+^ CD11b^+^ F4/80^-^ Ly6G^+^ (neutrophil) cells, and CD45^+^ CD11b^+^ F4/80^-^ Ly6G^-^ CD115^+^ (monocyte) cells in untreated B6 mice, anti-Ly6G and anti-rat κ immunoglobulin light chain antibody treated B6 mice, clodronate liposome treated B6 mice, and NSG mice. **c-e** Quantification of total number of (**c**) CD45^+^ CD11b^+^ (myeloid) cells, (**d**) macrophages, and (**e**) neutrophils infiltrating HA cryogels in untreated B6 mice, anti-Ly6G and anti-rat κ immunoglobulin light chain antibody treated B6 mice, clodronate liposome treated B6 mice, and NSG mice. (**f**) Schematic depicting workflow for in vivo Cy5-HA cryogel degradation study with gp91^phox-^ mice. (**g**) Representative IVIS fluorescence images of gel degradation and quantification by measuring total radiant efficiency normalized to initial day 3 timepoint of gp91^phox-^ mice and B6 mice. IVIS Images are on the same scale and analyzed using Living Image Software. Data in **c**, **d**, **e** represents mean ± s.d. of n=7-10 Cy5-HA cryogels, are representative of at least two separate experiments and compared using student’s t-test. Data in **g** represents mean ± s.e.m. of n=4-5 and were compared using two-way ANOVA with Bonferroni’s multiple comparison test. Parts of figures **3a and f** were created with BioRender.com.

Infiltration of total CD45^+^CD11b^+^ (myeloid) cells into HA cryogels of untreated B6 mice and T cell depleted B6 mice were similar after 1- and 10-days post-injection (**Supplementary Fig. 5d**). While there were comparable myeloid cells in HA cryogels retrieved 1-day post-injection from B cell depleted B6 mice, by day 10 the number was about 66% lower than in untreated B6 mice (**Supplementary Fig. 5e**). Similarly, myeloid cell infiltration in Cy5-HA cryogels retrieved 1-day post-injection from macrophage depleted mice was unaffected, however by day ten the number of infiltrating myeloid cells was about 58% lower than untreated B6 mice (**Fig. 3c**). Neutrophil depletion in B6 mice reduced the total number of myeloid cells in Cy5-HA cryogels compared with the untreated B6 mice by about 77% 1-day and 68% 10-days post-injection respectively (**Fig. 3c**). In NSG mice myeloid cell infiltration in HA cryogels was 51% lower 1-day and 91% lower 10-days post-injection as compared to untreated B6 mice (**Fig. 3c**).

CD45^+^CD11b^+^F4/80^+^ (macrophage) infiltration in Cy5-HA cryogels retrieved from all groups except NSG mice reduced from 1- to 10-days (**Fig. 3d**, **Supplementary Fig. 5e**). Intraperitoneal (i.p.) administration of clodronate liposomes minimally affected macrophage infiltration in Cy5-HA cryogels even though it was effective in depleting peripheral blood monocytes (**Fig. 3d, Supplementary Fig. 3g, h**). In NSG mice, macrophage infiltration was 74% lower than in the untreated B6 mice on day 1 and remained unchanged 10-days post-injection (**Fig. 3d**).

In CD45^+^CD11b^+^F4/80^-^Ly6G^+^ (neutrophil) depleted B6 mice, an additional intracellular Ly6G staining step was included, as the method of neutrophil depletion is known to induce internalization of the Ly6G receptor (**Supplementary Fig. 5f**) ^28^. Neutrophil infiltration in Cy5-HA cryogels retrieved from untreated B6 and T cell depleted B6 mice was comparable between 1- to 10-days post-injection (**Fig. 3e**, **Supplementary Fig. 5g**). B cell depletion did not affect the initial neutrophil infiltration 1-day post-injection compared with untreated controls but reduced the number of infiltrating neutrophils by 10-days post-injection (**Supplementary Fig. 5g**). As expected, neutrophil depletion significantly reduced initial neutrophil infiltration, by about 97%, compared to the untreated control. In this group, neutrophils constituted less than 50% of infiltrating myeloid cells at all timepoints assessed. Despite an increase by day 10, attributable to the internalization of the Ly6G receptor which led to an approximate 4-fold increase in the infiltrating neutrophil fraction (**Supplementary Fig. 5f**), the number of infiltrating neutrophils were still 84% lower compared with untreated B6 mice (**Fig. 3e**). Macrophage depletion did not affect the initial neutrophil infiltration 1-day post-injection compared with untreated B6 mice but reduced the number of infiltrating neutrophils by 65% compared with untreated B6 mice by day 10. HA cryogels retrieved from NSG mice had 50% fewer neutrophils than those from untreated B6 mice on day 1 and very few to none were found by day 10 post-injection (**Fig. 3e**). In NSG mice neutrophils constituted over 90% of total myeloid cells on day 1 but decreased to about 8% by day 10 (**Fig. 3e**). This observation along with minimal Cy5-HA cryogel degradation in NSG mice (**Fig. 2d**), supported a key role of functional neutrophils in mediating degradation. In all groups, the infiltration of CD45^+^CD11b^+^F4/80^-^ Ly6G^-^CD115^+^ (monocyte) cells were minimal and constituted a negligible portion of total infiltrating myeloid cells (**Supplementary Fig. 5h, i**).

To provide additional confirmation of infiltrating neutrophils and macrophages, we used immunohistochemical (IHC) staining to assess for Ly6G^+^ and F4/80^+^ cells respectively in untreated B6 mice and NSG mice at 1-, 5-, and 10-days post-injection. Staining on day 1 corroborated the flow cytometry data in that there were more neutrophils than macrophages within the Cy5-HA cryogels (**Supplementary Fig. 5i, Supplementary Fig. 6a**). On subsequent days, non-specific debris precluded accurate assessment in Cy5-HA cryogels retrieved from B6 mice (**Supplementary Fig. 6a**). As a result of non-specific staining of debris at later timepoints, IHC was only conducted on Cy5-HA cryogels excised 1-day after injection in macrophage depleted, neutrophil depleted, T cell depleted, and B cell depleted mice. Staining of these samples confirmed the presence of both Ly6G and F4/80 in Cy5-HA cryogels confirming flow cytometry data (**Supplementary Fig. 5i, Supplementary Fig. 6b**). In NSG mice, IHC staining of Ly6G^+^ and F4/80^+^ cells followed the results from flow cytometry analysis. Significantly more neutrophils than macrophages in Cy5-HA cryogels were observed 1-day after injection (**Supplementary Fig. 6c**). On day 5, there were significant macrophage and neutrophil infiltrates (**Supplementary Fig. 6c**) and by day 10, the neutrophil infiltration reduced significantly as expected from flow cytometry analysis (**Supplementary Fig. 5h, Supplementary Fig. 6c**).

To further characterize the role of functional neutrophils, we compared the degradation of Cy5-HA cryogels in B6 and B6.129S-Cybb^tm1Din^ (gp91^phox-^) mice. Affected hemizygous male gp91^phox-^ mice have a defect in the NADPH oxidase enzyme, which renders mice deficient in neutrophil function through the production of reactive oxygen species ^29, 30^. Cy5-HA cryogels were injected in gp91^phox-^ mice and B6 mice and degradation was quantified using IVIS (**Fig. 3f**). Cy5-HA cryogels did not degrade appreciably in the gp91^phox-^ over the course of the two-month study whereas the Cy5-HA cryogels in B6 mice degraded within 4 weeks, as expected (**Fig. 3g**).

Taken together, these results suggest that inducing immune deficiency by depletion affects cell infiltration in Cy5-HA cryogels but does not affect degradation. However, deficiencies which functionally impair neutrophils, modeled by NSG and gp91^phox-^ mice are sufficient to significantly affect Cy5-HA cryogel degradation.

### HA cryogels are neutrophil responsive in post-HSCT mice

We next quantified Cy5-HA cryogel degradation in post-HSCT mice. B6 recipients were irradiated 48 hours prior to i.v. injection of lineage depleted hematopoietic stem cells (2 x 10^5^ cells, ∼87% depleted) isolated from bone marrow of syngeneic B6 donor mice (**Supplementary Fig. 7a**). Concurrently, B6 recipients and control mice (B6, non-irradiated that do not receive a transplant) were injected subcutaneously with Cy5-HA cryogels, and the degradation rate was compared (**Fig. 4a**). In contrast to non-irradiated mice, a steady fluorescence signal was quantified for about 20 days in post-HSCT mice after which it decreased, corresponding to HA cryogel degradation, at a rate comparable to that in non-irradiated mice. The time interval to 50% of the initial fluorescence intensity was approximately 30 days in post-HSCT mice whereas in non-irradiated mice, a comparable decrease was achieved by day 13 (**Fig. 4b, Supplementary Fig. 7b**).

**Fig. 4.**
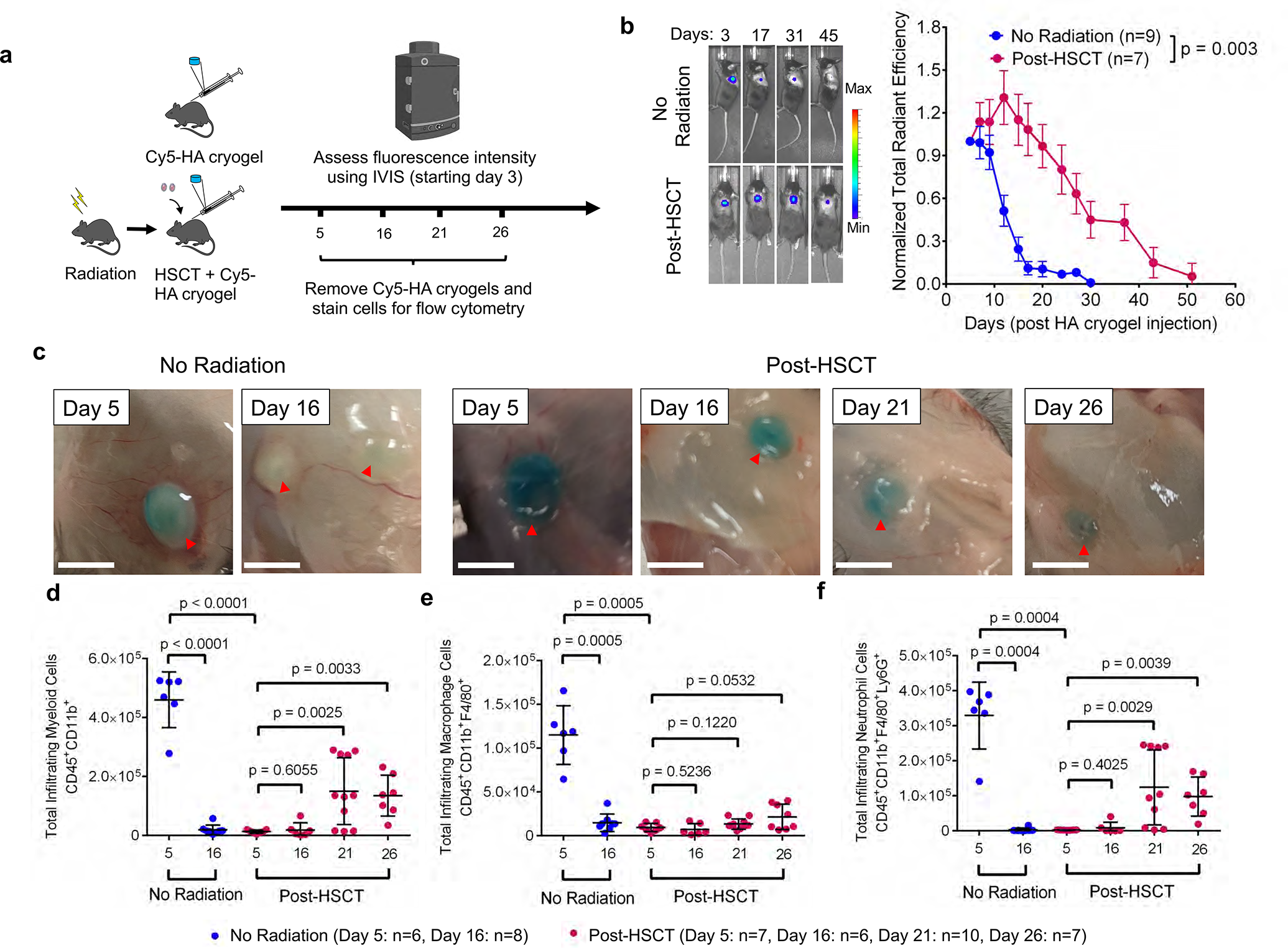
Degradation kinetics of HA cryogels is impaired during transient immunodeficiency following HSCT (a) Schematic depicting workflow for quantification of Cy5-HA cryogel degradation and innate immune cell infiltration in control (non-irradiated mice that do not receive a transplant) and post-HSCT B6 mice. (b) Representative IVIS fluorescence images of gel degradation in non-irradiated and post-HSCT mice. Tracking gel degradation by quantification of total radiant efficiency normalized to initial day 3 timepoint. (c) Photograph of Cy5-HA cryogels in non-irradiated mice 5- and 16-days post-injection and post-HSCT mice on days 5, 16, 21, and 26. **d-f** Cell infiltration of (**d**) CD45^+^CD11b^+^ (myeloid) cells, (**e**) CD45^+^CD11b^+^ F4/80^+^ (macrophage) cells, and (**f**) CD45^+^ CD11b^+^ F4/80^-^ Ly6G^+^ (neutrophil) cells into HA cryogels in non-irradiated mice 5- and 16-days post-injection and 5-, 16-, 21-, and 26- days post-HSCT. Data in **b** represents mean ± s.e.m. of n = 7-9 mice and is representative of at least two separate experiments. Data in **d**, **e**, **f** represents mean ± s.d. of n = 6-10 HA cryogels and are representative of at least two separate experiments. Data in **b** were compared using two-way ANOVA with Bonferroni’s multiple comparison test. Data in **d**, **e**, **f** were compared using student’s t-test. Part of figure **4a** was created with BioRender.com.

To quantify infiltrating myeloid subsets, Cy5-HA cryogels were excised on days 5 and 16 post-injection in non-irradiated mice and excised on days 5, 16, 21, and 26 in post-HSCT mice (**Fig. 4c**). Viability of infiltrating cells, quantified by negative AnnexinV staining, was initially lower 5- and 16-days post-injection, and increased by day 21 (**Supplementary Fig. 7c**). In non-irradiated B6 mice, the number of infiltrating myeloid cells decreased by 96% from days 5 to 16 post-injection (**Fig. 4d**), mirroring near-complete Cy5-HA cryogel degradation (**Fig. 4b, Supplementary Fig. 7b**). In contrast, myeloid cell infiltration in Cy5-HA cryogels was significantly delayed and 97% lower than that of the non-irradiated group, 5 days post-injection. In post-HSCT mice, appreciable myeloid infiltration was not quantified until about day 21 post-HSCT, which was still 67% lower when compared with HA cryogels from non-irradiated mice on day 5 (**Fig. 4d**).

Macrophage infiltration in Cy5-HA cryogels in non-irradiated mice decreased 87% from days 5 to 16 (**Fig. 4e**). On day 5 in post-HSCT mice, macrophage infiltration in Cy5-HA cryogels was reduced by about 92% compared to non-irradiated mice. By day 26, infiltrating macrophages in some post-HSCT mice were quantified but remained significantly lower than macrophage infiltration on day 5 in non-irradiated mice (**Fig. 4e**).

Neutrophils constituted a majority of the myeloid cells in Cy5-HA cryogels in non-irradiated mice on day 5, but not by day 16 (**Fig. 4f**) when the majority of myeloid cells were macrophages (**Supplementary Fig. 7d**). In contrast, very few cells were in HA cryogels retrieved on day 5 in post-HSCT mice with a near absence of neutrophils, in contrast with non-irradiated mice at the same timepoint. Neutrophil infiltration in Cy5-HA cryogels was quantified 21 days post-injection but was still 62% lower than on day 5 in non-irradiated mice (**Fig. 4f**). In post-HSCT mice, macrophages constituted most of the cell infiltrates 5-and 16-days after injection, whereas a majority of myeloid cells were neutrophils on days 21 and 26 (**Supplementary Fig. 7d**). This data supports that irradiation reduces myeloid infiltration in Cy5-HA cryogels, delays cryogel degradation, and degradation coincides with neutrophil recovery.

To assess whether the uniqueness of the results could be attributed to the HA cryogels, we compared the results with hydrolytically degradable oxidized alginate (OxAlg), also functionalized with Tz and Nb (**Supplementary Fig. 7e**). Unlike HA, OxAlg is not a substrate for endogenous enzymes ^31, 32^. Tz-functionalized OxAlg was functionalized with Cy5 and Cy5-OxAlg cryogels were formed in the same manner as high-DOS Cy5-HA cryogels. In vitro, Cy5-OxAlg cryogels fully degraded in 1x PBS over 9-days (**Supplementary Fig. 7f**). In contrast to Cy5-HA cryogels, Cy5-OxAlg cryogels injected in B6 and post-HSCT B6 mice degraded rapidly at a comparable rate, with approximately 70% reduction in fluorescence signal within 24 hours post-injection (**Supplementary Fig. 7g**).

### HA cryogels sustain G-CSF delivery and enhance post-HSCT reconstitution of neutrophils

We sought to leverage the delay in post-HSCT degradation of HA cryogels to mediate G-CSF release and enhance neutrophil recovery. As frequent bleeding to measure serum G-CSF concentrations is challenging in post-HSCT mice, we assessed G-CSF release from HA cryogels by labeling G-CSF with Cy5 and measuring the signal at the site of HA cryogel injection using IVIS microscopy. 1μg of Cy5-labeled G-CSF (Cy5 G-CSF) was encapsulated in HA cryogels and one cryogel was injected either in 1-day post-HSCT or in non-irradiated B6 mice. Encapsulated Cy5 G-CSF was quantified using IVIS and normalized to the initial 8-hour timepoint fluorescence signal (**Fig. 5a**). Cy5 G-CSF release, assessed by fluorescence attenuation, from non-irradiated mice proceeded in a sustained manner immediately post-injection with over 80% released after approximately 12-days post-injection. In post-HSCT mice, 20% Cy5 G-CSF released after approximately 12-days post-injection and subsequently released in a sustained manner (**Fig. 5b**). The time to 50% fluorescence intensity in non-irradiated mice was 5.9 ± 3.0 days compared to 15.5 ± 5.9 days in post-HSCT mice (**Fig. 5c**). To approximate G-CSF pharmacokinetics (PK) in the blood, the release profile of G-CSF from HA cryogels in post-HSCT mice was modeled as a piecewise function (**Supplementary Note 2**). We then sought to assess the effect of G-CSF delivery on peripheral blood neutrophil recovery and acceleration of Cy5-HA cryogel degradation. We compared mice receiving either two blank Cy5-HA cryogels or two G-CSF-encapsulated HA-cryogels and, as a positive control, we included mice with blank Cy5-HA cryogels injected systemically with 2μg pegylated (PEG) G-CSF (**Fig. 5d**), corresponding to the clinical-equivalent dose for mice ^33, 34^. Mice were bled at pre-determined timepoints, and peripheral blood neutrophil concentration, quantified by flow cytometry, was consistently higher when G-CSF from Cy5-HA cryogels was delivered, and comparable with PEG G-CSF treatment than in mice which received blank Cy5-HA cryogels (**Fig. 5e**). Moreover, Cy5-HA cryogel degradation was accelerated with G-CSF or PEG G-CSF treatment (**Fig. 5f**). These results support that G-CSF release from HA cryogels can improve neutrophil recovery in lethally radiated mice and Cy5-HA cryogel degradation may simultaneously be used as an indicator of functional neutrophil recovery.

**Fig. 5.**
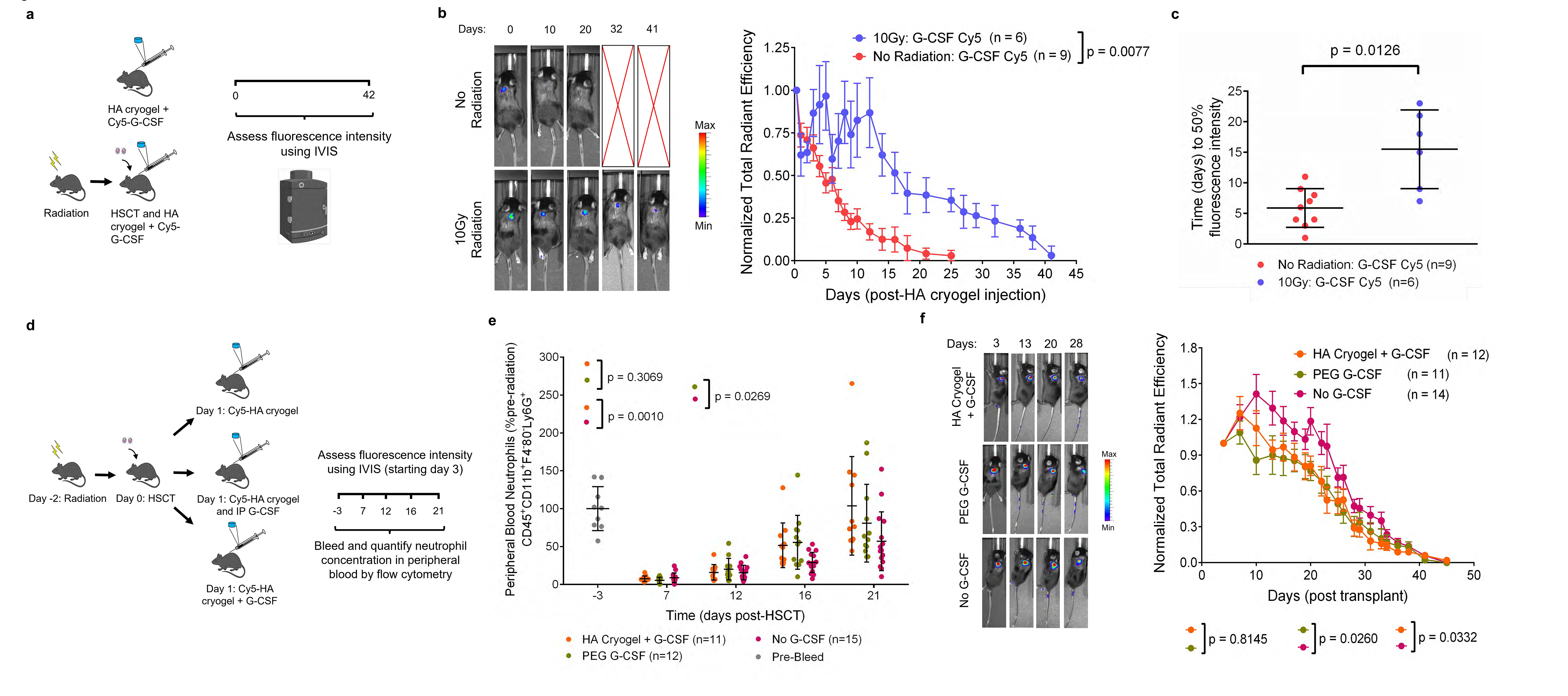
Enhanced reconstitution of peripheral blood neutrophil cells (a) Schematic depicting outline of study to quantify Cy5 G-CSF release from HA cryogels in non-irradiated, non-transplanted B6 mice and post-HSCT B6 mice. (b) Representative IVIS fluorescence images of Cy5 G-CSF release from HA cryogels and quantification by measuring total radiant efficiency normalized to initial 8-hour timepoint. IVIS Images are on the same scale and analyzed using Living Image Software. (c) Time to 50% fluorescence intensity for Cy5 G-CSF encapsulated within HA cryogels in non-irradiated and post-HSCT mice. (d) Schematic depicting outline of study to quantify neutrophil reconstitution rate and Cy5-HA cryogel degradation rate in post-HSCT mice using G-CSF encapsulated Cy5-HA cryogels. (e) Peripheral blood reconstitution of neutrophils in post-HSCT mice, normalized to pre-irradiation neutrophil counts from a random subset of mice. (**f**) Representative in vivo imaging system (IVIS) fluorescence images of gel degradation in mice and measuring Cy5-HA cryogel degradation in vivo by quantification of total radiant efficiency normalized to initial day 3 timepoint. IVIS Images are on the same scale and analyzed using Living Image Software. Data in **b** represents mean ± s.e.m. of n = 6-9 mice. Data in **c** represents mean ± s.d. of n = 6 – 9 mice. Data in **e** represents mean ± s.d. of n = 11-15 mice and is representative of at least two separate experiments. Data in **f** represents ± s.e.m. of n = 11-14 Cy5-HA cryogels and is representative of at least two separate experiments. Data in **b, f** were compared using two-way ANOVA with Bonferroni’s multiple comparison test. Data in **c** were compared using student’s t-test. Data in **e** were compared using mixed-effect regression model with random intercepts. Parts of figures **5a and d** were created with BioRender.com.

## Discussion

Here we demonstrate that an immune responsive biodegradable HA cryogel scaffold provides sustained G-CSF release and accelerates post-HSCT neutrophil recovery in mice which, in turn, accelerates HA cryogel degradation in vivo. Harnessing post-HSCT immune deficiency to sustain G-CSF release is distinct conceptually from other methods of drug delivery. It is well established that immune cells sense implanted materials as non-self and mount a well-characterized sequential response to isolate the implant in a fibrous capsule ^35–37^. In this work, we observed neutrophil infiltration during the acute stages of inflammation and show them to be key mediators in HA cryogel degradation. Our finding is consistent with prior reports that have supported neutrophils as key mediators of shaping the early implant microenvironment and for in vivo destruction of implanted polymeric materials by neutrophil-derived oxidants ^38–40^. The finding of primarily myeloid-lineage immune cell populations within the HA cryogel is consistent with previous observations of cell infiltration occurring within scaffolds of a similar composition ^41, 42^. We demonstrate that the encapsulation and release of G-CSF from the polymer scaffold mediated recovery of neutrophils in the peripheral blood, significantly faster than control mice receiving blank HA cryogels and comparable to pegylated GCSF, which accelerated HA cryogel degradation.

HA was selected as the primary constituent polymer as it is ubiquitous in the extracellular matrix and has a long history of clinical use as a biodegradable material in a range of biomedical applications ^43–48^. In this work, commercially purchased HA was derivatized with bioorthogonal Tz and Nb groups to facilitate crosslinking without the need for external energy input or addition of external agents such as stabilizers and catalysts ^49, 50^, which can make it challenging to purify the final product. The use of HA-Tz and HA-Nb also facilitated cryogelation at a slower rate, compared to free-radical polymerization methods, and consequently provided enhanced control over the crosslinking process ^51, 52^. Moreover, other common cross-linking strategies that directly target the carboxylic acid or hydroxyl side chains groups and unreacted agents may inadvertently react with encapsulated proteins ^53–56^. Further, Tz can be quantified spectroscopically and the DOS was readily assessed ^57, 58^. Consistent with prior work, our results show that DOS affected the rate of HA cryogel degradation by enzymatic cleavage ^22, 62^. On the other hand, the paradoxical observation that DOS did not affect in vivo degradation is also consistent with prior work that has demonstrated that partial degradation of HA by non-enzymatic means in vivo overcomes steric factors which might otherwise hinder enzymatic access to HA and, in our work, facilitated equalization of the in vivo degradation rate of low-and high-DOS HA cryogels ^59^.

Our results support that activated neutrophils mediate degradation of HA cryogels in vivo, consistent with past reports of the role of reactive oxygen species from activated neutrophils in mediating HA degradation ^60–63^ and of neutrophils in the acute phase of the foreign body response ^35, 36^, which further clarifies how immune deficiency impacts the rate of degradation ^9, 64^. We found that while despite successfully depleting neutrophils in the peripheral blood, antibody-based depletion did not achieve a similar depletion of infiltrating neutrophils in HA cryogels and degradation was unaffected in B6 mice. In NSG mice, which have defective adaptive and innate immune cells, Cy5-HA cryogels degraded minimally over 3 months and neutrophil infiltration into the HA cryogel was not sustained. The observation is consistent with the well-documented lack of adaptive immune cells, impaired innate immune cell subsets (e.g. macrophages) and a lack of a functional complement system which affects the activation of neutrophils in these mice ^65–68^. We expanded upon these results by quantifying Cy5-HA cryogel degradation in gp91^phox-^ mice, which are on the B6 background, but gp91^phox-^ neutrophils in affected hemizygous male mice lack superoxide production ^29, 30^. The functional deficiency of neutrophils in gp91^phox-^ is similar to the clinical observations of defective respiratory burst and phagocytosis affecting neutrophils in chronic granulomatous disease, in which there are normal neutrophil counts but impaired oxidative killing ^29^. In these mice, the absence of appreciable degradation of Cy5-HA cryogels provides additional support for the key role of functional neutrophils in facilitating degradation.

The key role of functional neutrophils in HA cryogel degradation was further validated in post-HSCT mice, modeling transient innate immune deficiency. Unlike antibody-based depletion, irradiating mice achieves elimination of all immune cells and neutrophils predominate the earliest immune cells that reconstitute post-HSCT ^69^. Similar to gp91^phox-^ mice, post-HSCT respiratory burst and phagocytosis of neutrophils are generally decreased in humans for up to 3 months, underscoring the importance of qualitatively assessing functionality of neutrophils ^70^. We found that Cy5-HA cryogel degradation was delayed until neutrophil infiltration into Cy5-HA cryogels recovered, further supporting the role of neutrophils in mediating HA degradation and the immune responsive behavior of HA cryogels. These results are consistent with past reports in which rapid neutrophil infiltration and activation have been identified as one of the earliest cellular events of the foreign body response ^2, 71^. In contrast, we show that OxAlg cryogels, which have hydrolytically labile groups but are not a substrate for endogenous enzymes, degrade rapidly in vivo at similar rates in immune competent and post-HSCT mice ^31, 32, 72^. These observations characterizing the importance of neutrophils in HA cryogel degradation support the unique immune responsiveness of HA cryogels.

Similar to PEG-G-CSF, we demonstrate the effect of G-CSF release from HA cryogels is neutrophil-dependent ^73^, and therefore might be characterized as self-regulating. However, in contrast to PEG-G-CSF, G-CSF delivery from HA cryogels avoids the potential of pre-existing or induced anti-PEG antibody (APA)-mediated rapid clearance ^74, 75^. In immune competent mice, it has been demonstrated that the administration of PEG-G-CSF at a clinically-relevant single dose elicits anti-PEG IgM antibodies in a dose-dependent manner which subsequently accelerates clearance of a second PEG-G-CSF dose via an anti-PEG IgM-mediated complement activation ^21^. De novo anti-PEG antibody induction may not require T cell activation ^76^and therefore could also be induced in post-HSCT immunodeficient hosts. PEG G-CSF may therefore be less effective with pre-existing or induced APA ^77^.

Therapy-induced neutropenia substantially limits the applicability of therapies that could be life-saving. HA cryogels not only deliver G-CSF in a sustained manner to enhance neutrophil regeneration, while avoiding the potential of APA-mediated enhanced clearance, but also show a responsive degradation behavior. Collectively, our findings support that the HA cryogels might be leveraged to enhance and functionally assess neutrophil functionality and aid in treatment-related decisions for recipients of myelosuppressive therapy.

## Supplementary Note 1

The second order rate constant (k) for Tz-Nb reaction has been previously estimated to be 1.3 - 1.7 M^-1^s^-1^ at 21°C (room temperature).^1, 2^ In our system, for high-DOS Tz-HA and Nb-HA the concentration is 0.55mM at the start of the reaction. The reaction rate can be calculated as:

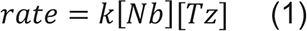

We calculated the rate of reaction to be about 40µM/s from equation 1 and the time to completion to be about 46.3 minutes at 21°C. In our system, the initial temperature is 4°C and therefore the actual time for completion of the reaction would be significantly longer.

As the solution cools and freezes during the crosslinking process, we estimated the freezing time. First, we determined the energy required to freeze 30uL of HA solution starting from 4°C and ending at 0°C using:

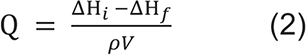

The energy required to freeze 30µL HA solution from 4°C to 0°C is 10.5J^3^. Since the HA solution is very dilute (0.6 wt%), we have approximated the enthalpy of formation and density to that of water.

To calculate the freezing time, we need to estimate the rate of energy extraction from the HA solution. Since the teflon cryomold is pre-cooled to -20°C and rests on a metal shelf in the freezer, we can estimate the rate of freezing using the thermal conductivity of teflon, thickness of teflon (25mm), and conductive heat transfer area (5.75mm) using equation 3. To simplify the analysis, we assume that convective heat loss at air-cryogel interface is negligible.

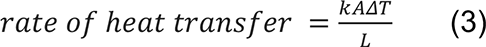

The rate of heat transfer is calculated to be 0.010J/s and therefore the time to reach 0°C is ∼16.8 minutes. Since we are ignoring conductive heat loss through the edge of the cryogels and convective heat loss through the top, the calculated time represents an overestimation for the freezing time but is still significantly below that of the time to reaction completion. We have also experimentally verified that a 30uL drop of Tz-HA/Nb-HA solution freezes in about 10 minutes.

## Supplementary Note 2

To model G-CSF pharmacokinetics (PK) in the blood, the release profile of G-CSF from HA cryogels in post-HSCT mice was modeled as a piecewise function (**Fig. SN1**). Phase 1 was used to estimate G-CSF release from days 0-12 and phase 2 was used for days 12-40. We adopted parameters from previous reports^1, 2^.

**Figure SN1.**
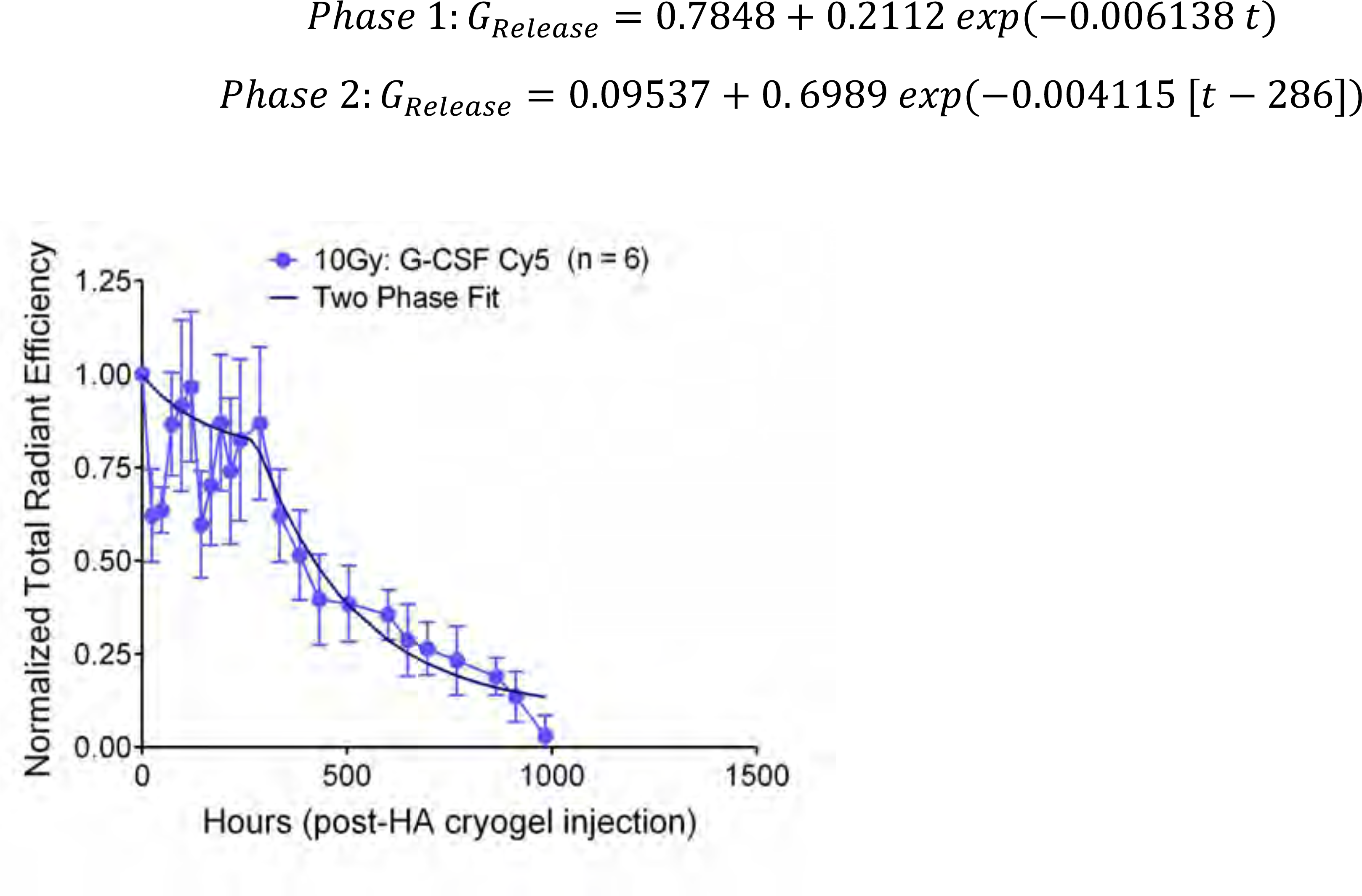
Two-phase curve fit of G-CSF release from HA cryogel post-HSCT (as depicted in Fig. 5b).

The model takes into account endogenous production of G-CSF and assumes two HA cryogels loaded with 1µg of G-CSF each as sources. Renal clearance and internalization by neutrophil progenitors are consumption terms (**Equations 1,2** and **Fig. SN2a**).

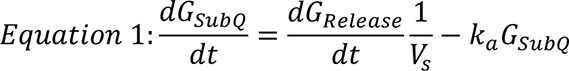

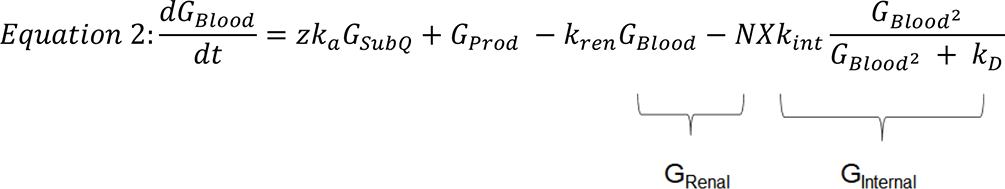

**Figure SN2.**
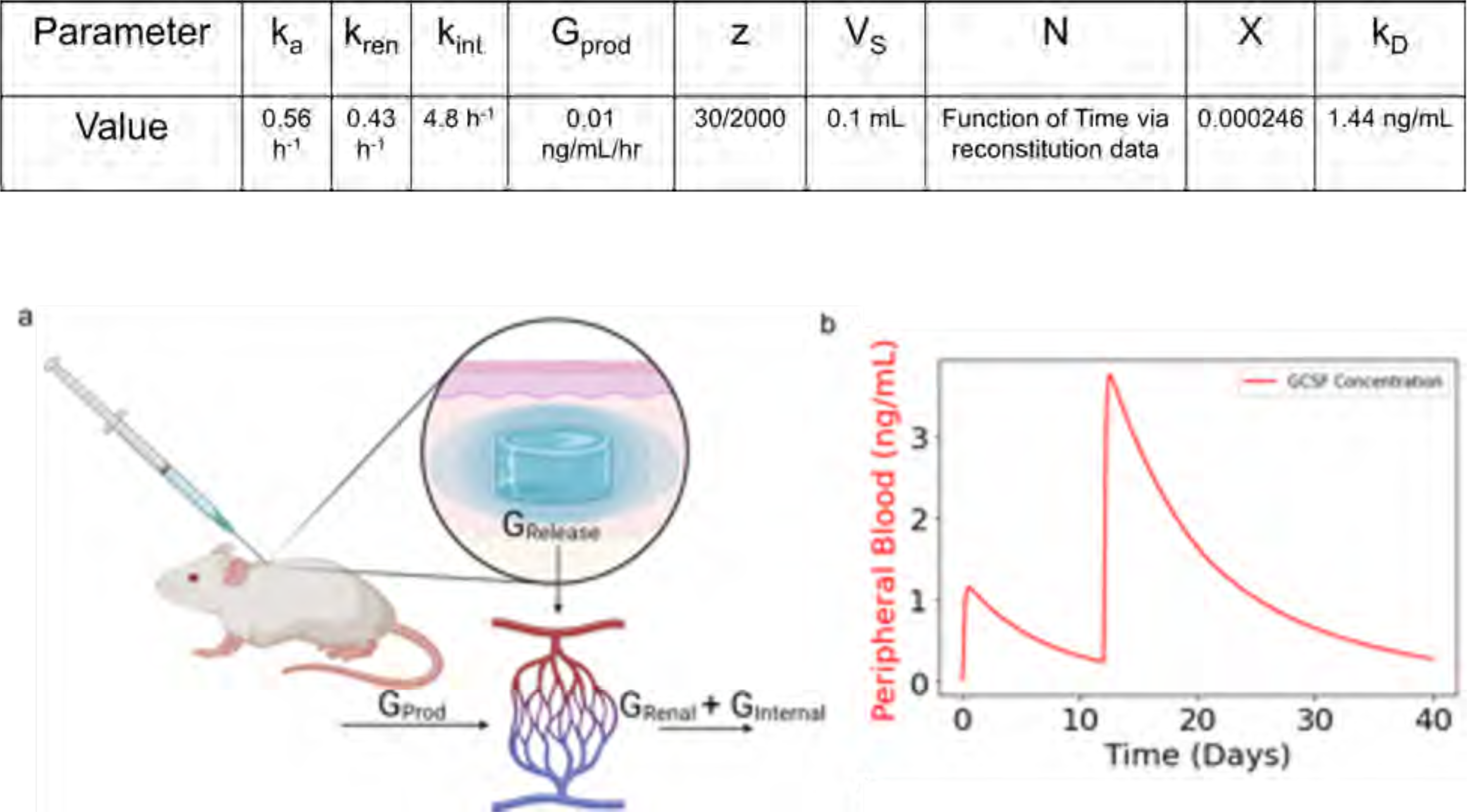
(a) Schematic depicting sources of G-CSF production and clearance in peripheral blood. (b) Predicted G-CSF concentration in peripheral blood from two subcutaneously administered G-CSF encapsulated HA cryogels based on experimentally determined G-CSF release from HA cryogels.

The resulting G-CSF concentration in the peripheral blood of mice is estimated in **Fig. SN2b**. The code underlying the model can be found here: https://github.com/Shah-Lab-UCSD/GCSF-Release-Mod

## Methods

### General methods and statistics

Sample sizes for animal studies were based on prior work without use of additional statistical estimations. Results were analyzed where indicated using student’s t-test and two-way ANOVA with Bonferroni’s test using Graphpad Prism software. Mixed-model linear regression was conducted using IBM SPSS statistical package. Alphanumeric coding was used in blinding for pathology samples and cell counting.

### Chemicals

Sodium hyaluronate (MW 1.5-2.2 MDa, Pharma Grade 150, lot: 18011K) and sodium alginate (MW ∼250 kDa, Pronova UP MVG) were purchased from NovaMatrix. (2-morpholinoethanesulfonic acid (MES), sodium chloride (NaCl), sodium hydroxide (NaOH), *N*-hydroxysuccinimide (NHS), 1-ethyl-3-(3-dimethylaminopropyl)-carbodiimide hydrochloride (EDC), sodium periodate (311448) and ammonia borane (AB) complex (682098) were purchased from Sigma-Aldrich. (4-(1,2,4,5-tetrzain-3-yl)phenyl)methanamine (tetrazine amine) was purchased from Kerafast (FCC659, lot: 2014). 1-bicyclo[2.2.1]hept-5-en-2-ylmethanamine (norbornene amine) was purchased from Matrix Scientific (# 038023, lot: M15S). Cy5-tetrazine amine was purchased from Lumiprobe (lot: 9D2FH). 1kDa molecular weight cutoff (MWCO) mPES membrane was purchased from Spectrum (S02-E001-05-N).

### Derivatization of HA

Tetrazine functionalized HA (HA-Tz) or norbornene functionalized HA (HA-Nb) were prepared by reacting tetrazine amine or norbornene amine to HA using EDC/NHS carbodiimide chemistry. Sodium hyaluronate was dissolved in a buffer solution (0.75% wt/vol, pH ∼ 6.5) of 100mM MES buffer. NHS and EDC were added to the mixture to activate the carboxylic acid groups on the HA backbone followed by either tetrazine amine or norbornene amine. HA was assumed to be 1.8 MDa for purposes of conjugation reactions. To synthesize 7% DOS HA-Tz (high-DOS), the molar ratios of HA:EDC:NHS:tetrazine are 1:25000:25000:2500. To synthesize 0.8% DOS HA-Tz (low-DOS), the molar ratios of HA:EDC:NHS:tetrazine are 1:2860:2860:286. To synthesize HA-Nb, the molar ratios of HA:EDC:NHS:norbornene are 1:25000:25000:2500. Each reaction was stirred at room temperature for 24 hours and transferred to a 12,000Da MW cutoff dialysis sack (Sigma Aldrich) and dialyzed in 4L of NaCl solutions of decreasing molarity (0.125M, 0.100M, 0.075M, 0.050M, 0.025M, 0M, 0M, 0M, 0M) for 8 hours per solution. After dialysis, solutions containing HA-Tz or HA-Nb were frozen overnight and lyophilized (Labconco Freezone 4.5) for 48 hours. Cy5 conjugated HA-Nb (Cy5-HA-Nb) was synthesized following a previously described technique with some modifications ^78^. 0.8mg of Cy5-Tz was reacted with 100mg of HA-Nb at 0.2 wt/vol in DI water for 24 hours at 37 °C and purified by dialysis in DI water using a 12,000Da MW cutoff dialysis sack for 48 hours. Dialysis water bath was changed once every 8 hours. Cy5-HA-Nb was then frozen overnight and lyophilized for 48 hours.

### Preparation of oxidized alginate

Alginate was oxidized by mixing a 1% wt/vol solution of sodium alginate in DI water with an aqueous solution of 23 mM sodium periodate (Sigma Aldrich) to achieve a 1:586 molar ratio of alginate: periodate. The reaction was stirred in the dark at room temperature overnight. Sodium chloride (1.8 grams/gram of alginate) was added to solution to achieve a 0.3 M solution, followed by purification via tangential flow filtration (TFF) using a mPES 1kDa molecular weight cutoff (MWCO) membrane (Spectrum) and sequential solvent exchanges with 0.15 M – 0.10 M – 0.05 M and 0.0 M sodium chloride in DI water. The resulting solution was treated with ammonia borane (AB) complex (Sigma Aldrich) at 1:4 alginate:AB molar ratio and stirred at room temperature overnight. Sodium chloride (1.8 grams/gram of alginate) was added to solution to achieve a 0.3 M solution, followed by purification via TFF using a 1kDa MWCO mPES membrane and sequential solvent exchanges with 0.15 M – 0.10 M – 0.05 M and 0.0 M sodium chloride in DI water. The resulting solution was lyophilized to dryness.

### Derivatization of oxidized alginate

To synthesize tetrazine and norbornene functionalized oxidized alginate (OxAlg-Tz, OxAlg-Nb respectively), oxidized alginate, prepared as described above, was solubilized in 0.1 M MES buffer,0.3 M sodium chloride, pH 6.5 at 1%wt/vol. NHS and EDC were added to the mixture followed by either tetrazine or norbornene. The molar ratio of oxidized alginate:NHS:EDC:tetrazine or norbornene was 1:5000:5000:1000. The reaction is stirred in the dark at room temperature overnight. The resulting solution is centrifuged at 4700 rpm for 15 minutes and filtered through a 0.2-micron filter. The solution is purified via TFF using a mPES 1kDa molecular weight cutoff (MWCO) membrane and sequential solvent exchanges with 0.15 M – 0.10 M – 0.05 M and 0.0 M sodium chloride in DI water. The purified solution is treated with activated charcoal (1 gram / gram of alginate) for 20 minutes at room temperature. The slurry is filtered through 0.2-micron filter and the filtrate is lyophilized to dryness.

### Endotoxin Testing

Endotoxin testing of high-DOS HA-Tz and Cy5-HA-Nb were conducted using a commercially available endotoxin testing kit (88282, Thermo Fisher Scientific, lot: VH310729) and following manufacturer’s instructions. High-DOS HA-Tz and Cy5-HA-Nb were solubilized at 0.6 wt% in endotoxin free water and samples were tested in technical triplicates. To calculate endotoxin content of a single HA cryogel, the EU/mL concentration for high-DOS HA-Tz and Cy5-HA-Nb were divided by 2 (relative concentrations of HA-Tz and HA-Nb in HA cryogels are 0.3 wt%) and multiplied by 0.03 (30uL of volume per HA cryogel). EU/kg was calculated based on 2 HA cryogels administered into a mouse with an average weight of 20 grams.

### Cryogel development

We followed a previously described cryogelation method^78–80^. To form cryogels, aqueous solutions of 0.6% wt/vol HA-Tz and HA-Nb or OxAlg-Tz and OxAlg-Nb were prepared by dissolving lyophilized polymers into deionized water and left on a rocker at room temperature for a minimum of 8 hours to allow for dissolution. The aqueous solutions were then pre-cooled to 4°C before cross-linking to slow reaction kinetics. HA-Tz and HA-Nb or OxAlg-Tz and OxAlg-Nb solutions were mixed at a 1:1 volume ratio, pipetted into 30μL Teflon molds which were pre-cooled to -20 °C, and quickly transferred to a -20 °C freezer to allow for overnight cryogelation. Synthesis of Cy5-HA or Cy5-OxAlg cryogels follows the same protocol as above, substituting Cy5-HA-Nb for HA-Nb or Cy5-OxAlg-Nb for OxAlg-Nb.

### Pore size analysis of HA cryogels

For scanning electron microscopy (SEM), frozen HA cryogels were lyophilized for 24 hours and in a petri dish. Lyophilized HA cryogels were adhered onto sample stubs using carbon tape and coated with iridium in a sputter coater. Samples were imaged using secondary electron detection on a FEI Quanta 250 field emission SEM in the Nano3 user facility at UC San Diego. Fluorescence images of Cy5-HA cryogels were acquired using a Leica SP8 All experiments were performed at the UC San Diego School of Medicine Microscopy Core. Pore size quantification of SEM images and relative distribution of pore sizes of confocal images was doing using FIJI image processing package ^81^.

### HA cryogel pore-interconnectedness analysis

Cy5-HA cryogels were synthesized with low- and high-DOS HA-Tz and incubated in 1mL of FITC-labeled 10µM diameter melamine resin micro particles (Sigma Aldrich) at 0.29mg/mL concentration on a rocker at room temperature overnight. Fluorescence images of Cy5-HA cryogels with FITC-labeled microparticles were acquired using a Leica SP8 confocal. Interconnectedness of the HA cryogels was determined by generating 3D renderings of confocal z-stacks using FIJI imaging processing package and assessing fluorescence intensity of both the Cy5 and FITC channels with depth starting from the top of the HA cryogel. To determine the effect of injection on pore interconnectedness, HA cryogels were injected through a 16G needle prior to incubation in FITC-labeled microparticle solution. All experiments were performed at the UC San Diego School of Medicine Microscopy Core.

### In vitro degradation of Cy5-HA cryogels

Cy5-HA cryogels synthesized with low- and high-DOS HA-Tz and placed into individual 1.5mL microcentrifuge tubes (Thermo Scientific) with 1mL of 100U/mL Hyaluronidase from sheep testes Type II (HYAL2, H2126, Sigma Aldrich, lot: SLBZ9984) in 1x PBS. Degradation studies were conducted in tissue culture incubators at 37 °C. Supernatant from samples were collected every 24 – 72 hours by centrifuging the samples at room temperature at 2,000G for 5 minutes and removing 0.9mL of supernatant. Cy5-HA cryogels were resuspended by adding 0.9mL of freshly made 100U/mL HYAL2 in 1x PBS. Fluorescence measurements were conducted using a Nanodrop 2000 Spectrophotometer (Thermo Fisher Scientific) and these values were normalized to sum of the fluorescence values over the course of the experiment. All experiments were performed at UC San Diego.

### In vitro degradation of Cy5-OxAlg Cryogels

Cy5-OxAlg cryogels were placed into individual 1.5mL microcentrifuge tube with 1mL of 1x PBS. Degradation studies were conducted in tissue culture incubators at 37°C. Supernatant from samples were collected every 24 – 72 hours by removing visible Cy5-OxAlg cryogel material with tweezers and transferring to new 1.5mL microcentrifuge tube with 1mL of 1x PBS. Fluorescence measurements were conducted using a Nanodrop 2000 Spectrophotometer and these values were normalized to sum of the fluorescence values over the course of the experiment. All experiments were performed at UC San Diego.

### In vivo mouse experiments

All animal work was conducted at the Moores Cancer Center vivarium at UC San Diego, except NSG mouse IVIS imaging experiments, which were conducted at the Harvard Biological Research Infrastructure vivarium at Harvard University and approved by the respective Institutional Animal Care and Use Committee (IACUC). All animal experiments followed the National Institutes of Health guidelines and relevant AALAC-approved procedures. Female C57BL/6J (B6, Jax # 000664) and NOD.Cg-*Prkdc^scid^ Il2rg^tm1Wjl^*/SzJ (NSG, Jax # 005557) mice were 6-8 weeks at the start of the experiments. Male B6.129S-Cybb^tm1Din^ (gp91^phox-^, Jax # 002365) mice were 6-8 weeks old at the start of experiments. All mice in each experiment were age matched and no randomization was performed. The pre-established criteria for animal omission were failure to inject the desired cell dose in transplanted mice and death due to transplant failure. Health concerns unrelated to the study (e.g. malocclusion) and known mouse-strain specific conditions that affected measurements (e.g. severe dermatitis and skin hyperpigmentation in B6 mice) were criteria for omission.

### Immune depletion in mice

Neutrophil depletion in B6 mice was achieved by following the previously established protocol ^28, 82^. Briefly, 25μL of anti-mouse Ly6G antibody (1A8, Bio X Cell, lot: 737719A2)) was administered i.p. every day for the first week. Concurrently, 50μL of anti-rat κ immunoglobulin light chain antibody (MAR 18.5, Bio X Cell, lot: 752020J2) was administered every other day starting on the second day of depletion. After one week, the dose of the anti-mouse Ly6G antibody was increased to 50μL. Macrophage depletion in B6 mice was induced by i.p. administration of 100uL of clodronate liposomes (Liposoma, lot: C44J0920) every 3-days. B cell lineage depletion in B6 mice was induced by i.p. administration of 400μg of anti-mouse B220/CD45R antibody (RA3.31/6.1, Bio X Cell, lot: 754420N1) once every 3-days. T cell lineage depletion in B6 mice was induced by i.p. administration of 400μg dose of anti-mouse CD4 antibody (GK1.5, Bio X Cell, lot: 728319M2) and 400μg dose of anti-mouse CD8α antibody (2.43, Bio X Cell, lot:732020F1) once every 3-days. For all lineage depletion models, mice received intraperitoneal injections of 0.1mL (400μg) of antibodies or 0.1mL of clodronate liposome solution 3 days before subcutaneous HA cryogel or Cy5-HA cryogel injection. Depletion started 3-days prior to Cy5-HA cryogel administration to mice and continued until complete cryogel degradation or until mice were euthanized and cryogels retrieved for analysis. All experiments were performed at the Moores Cancer Center vivarium at UC San Diego.

### Transplant models

Irradiations were performed with a Cesium-137 gamma-radiation source irradiator (J.L. Shepherd & Co.). Syngeneic HSCT (B6 recipients) consisted of 1 dose of 1,000 cGy + 1 x 10^5^ lineage-depleted bone marrow cells from syngeneic B6 donors. Bone marrow cells for transplantation (from donors) or analysis were harvested by crushing all limbs with a mortar and pestle, diluted in 1x PBS, filtering the tissue homogenate through a 70 μm mesh and preparing a single-cell suspension by passing the cells in the flowthrough once through a 20-gauge needle. Total cellularity was determined by counting cells using a hemacytometer. Bone marrow cells were depleted of immune cells (expressing CD3ε, CD45R/B220, Ter-119, CD11b, or Gr-1) by magnetic selection using a Mouse Hematopoietic Progenitor Cell Enrichment Set (BD Biosciences # 558451, lot: 0114777). To confirm depletion, we incubated cells with a mix of Pacific Blue-conjugated lineage specific antibodies (antibodies to CD3, NK1.1, Gr-1, CD11b, CD19, CD4 and CD8) and with Sca-1 and cKit-specific antibodies for surface staining and quantification of Lineage^-^ fraction of cells, which were ≥ 87% lineage depleted. Subsequently, cells were suspended in 100μL of sterile 1x PBS and administered to anesthetized mice via a single retroorbital injection. All experiments were performed at the Moores Cancer Animal Facility at UC San Diego Health. All flow cytometry experiments were performed using an Attune^®^ NxT Acoustic Focusing cytometer analyzer (A24858) at UC San Diego.

### Subcutaneous cryogel administration

While mice were anesthetized, a subset received a subcutaneous injection of HA cryogel or OxAlg cryogel, which was suspended in 200μL of sterile 1x PBS, into the dorsal flank by means of a 16G needle positioned approximately midway between the hind- and forelimbs. The site of injection was shaved and wiped with a sterile alcohol pad prior to gel injection.

### In vivo degradation

In vivo Cy5-HA cryogel degradation was performed with Cy5-HA cryogels synthesized with low- and high-DOS Tz-HA in untreated B6 mice, immune deficient B6 mice, NSG mice, and gp91^phox-^ mice. In vivo Cy5 OxAlg cryogel degradation was performed in non-irradiated, non-transplanted B6 mice and B6 mice post-HSCT. In all cases, cryogels were administered into the dorsal flank of an anesthetized mouse and the fluorescent intensity of the Cy5-HA cryogel was quantified using an IVIS spectrometer (PerkinElmer) at predetermined timepoints and analyzed using LivingImage software (PerkinElmer). At each timepoint, mice were anesthetized and the area around the subcutaneous cryogel was shaved to reduce fluorescence signal attenuation. Fluorescence radiant efficiency, the ratio of fluorescence emission to excitation, was measured longitudinally as a metric to quantify fluorescence from subcutaneous cryogels. These values were normalized to the measured signal on day 3. All experiments were performed at the Moores Cancer Microscopy Core Facility at UC San Diego Health, with the exception of NSG mouse in vivo degradation experiments, which were performed at the Harvard Biology Research Infrastructure vivarium using an IVIS spectrometer (PerkinElmer).

### Flow cytometry analysis

Anti-mouse antibodies to CD45 (30-F11, lot: B280746), CD11b (M1/70, lot: B322056), CD4 (RM4-5, lot: B240051), CD8α (53-6.7, lot: B266721), B220 (RA3-6B2, lot: B298555), Ly6-G/Gr-1 (1A8, lot: B259670), lineage cocktail (17A2/RB6-8C5/RA3-6B2/Ter-119/M1/70, lot: B266946), Ly-6A/E/Sca-1 (D7, lot: B249343), and CD117/cKit (2B8, lot: B272462) were purchased from Biolegend. Anti-mouse F4/80 (BM8, lot: 2229150) and was purchased from eBioscience. All cells were gated based on forward and side scatter characteristics to limit debris, including dead cells. AnnexinV (Biolegend, lot: B300974) stain was used to separate live and dead cells. Antibodies were diluted according to the manufacturer’s suggestions. Cells were gated based on fluorescence-minus-one controls, and the frequencies of cells staining positive for each marker were recorded. To quantify T cells, B cells, monocytes, and neutrophils in peripheral blood, blood was first collected from the tail vein of mice into EDTA coated tubes (BD). Samples then underwent lysis of red blood cells and were stained with appropriate antibodies corresponding to cell populations of interest. To quantify infiltrating immune cells within Cy5-HA cryogels, mice were sacrificed, cryogels removed, and HA cryogels crushed against a 70-micron filter screen before antibody staining. Absolute numbers of cells were calculated using flow cytometry frequency. Flow cytometry was analyzed using FlowJo (BD) software. All flow cytometry experiments were performed using a Attune^®^ NxT Acoustic Focusing cytometer analyzer (A24858) at UC San Diego.

### Histology

After euthanasia, HA cryogels were explanted and fixed in 4% paraformaldehyde (PFA) for 24 hours. The fixed HA cryogels were then transferred to 70% ethanol solution. Samples were routinely processed and sections (5μm) were stained and digitized using an Aperio AT2 Automated Digital Whole Slide Scanner by the Tissue Technology Shared Resource at the Moores Cancer Center at UC San Diego Health. Digital slides were rendered in QuPath and positive cell detection was used to quantify the total number of mononuclear cells within each image. Quantification of mononuclear cell density was determined for each histological section.

### Immunohistochemistry (IHC)

Paraffin embedded HA cryogel sections were baked at 60 °C for 1 hr. Tissues were then rehydrated through successive washes (3x xylene, 2x 100% ethanol, 2x 95% ethanol, 2x 70% ethanol, di-water). After rehydration, antigen retrieval was conducted using Unmasking solution (Citrate based, pH 6) (Vector Laboratories, H-3300) at 95 °C for 30 minutes. Staining was performed using Intellipath Automated IHC Stainer (Biocare). A peroxidase block, Bloxall (Vector Laboratories, SP-6000) was performed for 10 minutes, followed by 2x washes in 1x tris-buffered saline with 0.1% Tween 20 (TBST, Sigma Aldrich), and a blocking step using 3% Donkey Serum for 10 minutes. Samples were stained using anti-Ly6G primary antibody (Rabbit, Cell Signaling Technology, 87048S) at 1:100 concentration for 1 hr. Samples were washed twice in TBST and anti-rabbit HRP polymer (Cell IDX, 2HR-100) was added for 30 minutes. Samples were washed twice in TBST and DAB (brown) Chromogen (VWR, 95041-478) was added for 5 minutes. This was followed by 2x washes in di-water, 5-minute incubation with Mayer’s Hematoxylin (Sigma, 51275), 2x washes in TBST, and 1x wash in di-water. Mounting was performed using a xylene-based mountant. IHC was performed by the Tissue Technology Shared Resource at the Moores Cancer Center at UC San Diego Health.

### G-CSF encapsulation

To quantify G-CSF release from HA cryogels, recombinant human G-CSF (300-23, Peprotech, lots: 041777 and 041877) was reacted with sulfo-Cy5 NHS ester (13320, Lumiprobe, lot 7FM7C) at a 1:250:25 molar ratio of G-CSF:EDC:sulfo-Cy5 NHS ester in MES buffer to form Cy5 G-CSF. Unreacted EDC and sulfo-Cy5 NHS ester was removed by overnight dialysis on a 10kDa dialysis membrane. 1μg of Cy5 G-CSF was added to 0.6% wt/vol HA-Tz solution before mixing with HA-Nb and cryogelation as described above. To track Cy5-HA cryogel degradation in mice which received G-CSF loaded Cy5-HA cryogels, the same protocol is followed substituting G-CSF for Cy5 G-CSF and Cy5-HA-Nb for HA-Nb.

### Neutrophil reconstitution models

Mice were irradiated and administered an autologous HSCT as described above. PEG G-CSF (MBS355608, MyBioSource, lot: R15/2020J) or G-CSF was injected i.p. Cy5-HA cryogel encapsulating G-CSF was injected subcutaneously as described above, 24 hours post-HSCT. Mice were bled at predetermined timepoints and relevant immune subsets were stained for flow cytometry.

## Contributions

**Matthew D. Kerr:** Conceptualization, methodology, validation, formal analysis, investigation, resources, data curation, writing – original draft, writing – review & editing and visualization. **David A. McBride:** Methodology, investigation, writing – review and editing. **Wade T. Johnson:** Investigation and writing – review and editing. **Arun K. Chumber:** Investigation and writing – review and editing. **Alexander J. Najibi:** Investigation and writing – review and editing. **Bo Ri Seo.** Investigation and writing – review and editing. **Alexander G. Stafford:** Resources. **David T. Scadden:** Funding acquisition. **David J. Mooney:** Conceptualization, writing – review and editing, and funding acquisition. **Nisarg J. Shah:** Conceptualization, writing - original draft, writing – review and editing, supervision, project administration, funding acquisition. All authors reviewed the manuscript and data, provided input and approved the submission.

## Acknowledgements

The work was supported in part by the American Cancer Society (IRG-15-172-45-IRG), National Multiple Sclerosis Society (PP-1905-34013), the National Institutes of Health (R03DE031009, P30AR073761, and R01DE013349), and the Blavatnik Biomedical Accelerator and the Hansjörg Wyss Institute for Biologically Inspired Engineering at Harvard University. M.D.K. and D.A.M. received NIH training grant support through the NCI (T32CA153915) and NIAMS (T32AR064194 and F31AR079921) respectively. The authors acknowledge assistance by the Moores Cancer Center Tissue Technology Shared Resource at UC San Diego Health, the Microscopy Core at UC San Diego School of Medicine, and the Biostatistics Unit of the Clinical and Translational Research Institute at UC San Diego supported by the National Institutes of Health (P30CA23100, P30NS047101 and UL1TR001442 respectively). This work was performed in part at the San Diego Nanotechnology Infrastructure (SDNI) of UCSD, a member of the National Nanotechnology Coordinated Infrastructure (NNCI), which is supported by the National Science Foundation under Grant No. ECCS-1542148. The content is solely the responsibility of the authors and does not necessarily represent the official views of the National Institutes of Health or the National Science Foundation.

## Data availability

The data that support the findings of this study are available from the corresponding author upon reasonable request.

## Supplementary Figure Captions

**Fig. S1.**
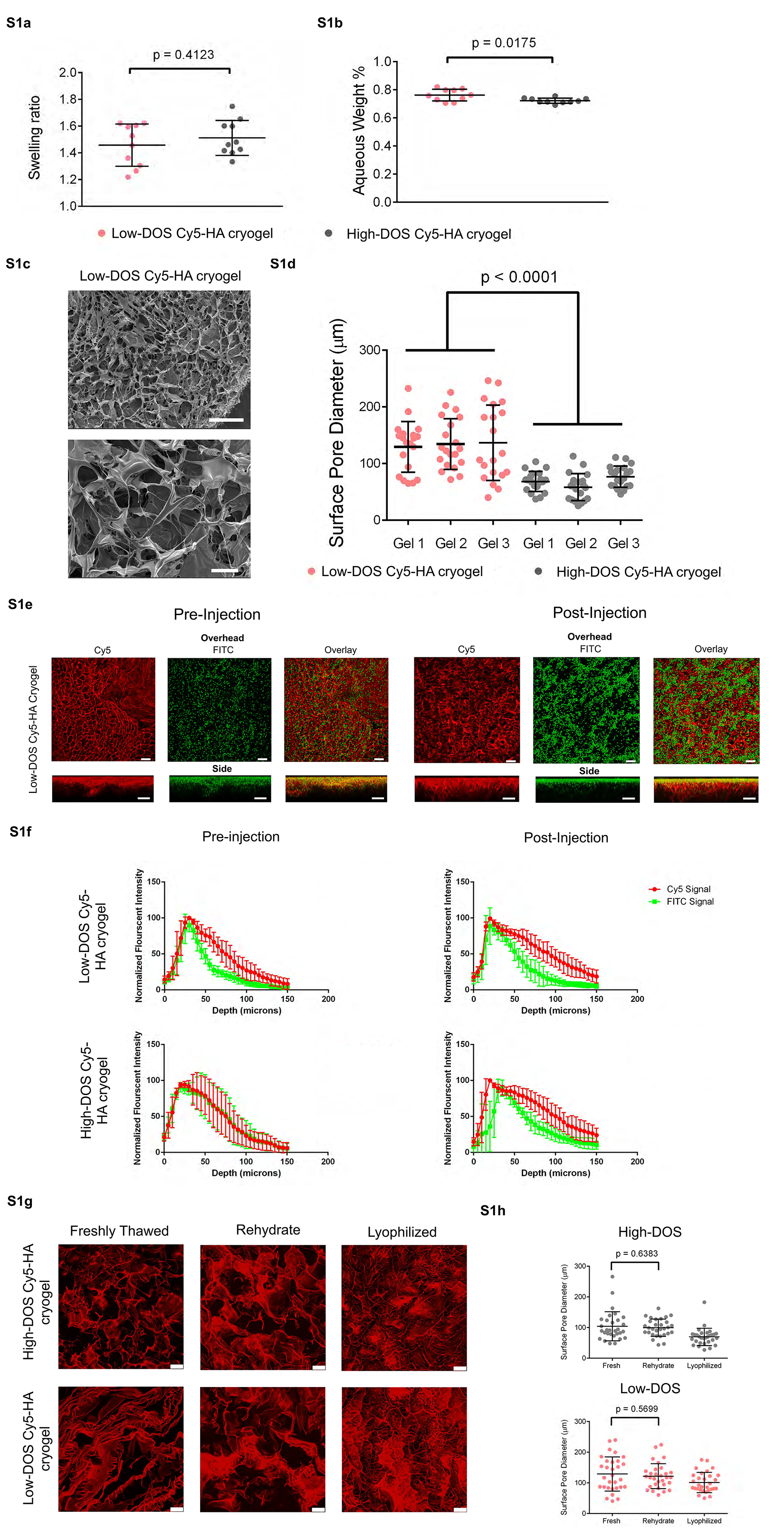
Supplementary HA cryogel materials characterization data (**a**) Volumetric swelling ratios for low- and high-DOS Cy5-HA cryogels. (**b**) Aqueous weight percentage of low- and high-DOS Cy5-HA cryogels. (**c**) Representative SEM image depicting low-DOS Cy5-HA cryogels. Top scale bar = 500μm, bottom scale bar = 100μm. (**d**) Average pore diameters of HA cryogels made from low-and high-DOS Cy5-HA cryogels measured from SEM images (20 measurements/cryogel, n = 3 each for low-and high-DOS HA cryogels). (**e**) Confocal microscopy images, overhead and side views, depicting low-DOS Cy5-HA cryogels both pre-injection and post-injection incubated with 10μm FITC-labeled microparticles. Scale bar = 100μm. (**f**) Quantification of confocal images showing penetration of 10μm FITC-labeled microparticles into both low- and high-DOS Cy5-HA cryogels pre-and post-injection. (**g**) Representative confocal microscopy images of low- and high-DOS Cy5-HA cryogels after thawing, after lyophilization, and after lyophilization and rehydration. (**h**) Average surface pore diameter of Cy5-HA cryogels measured from confocal images. Data in **a** and **b** represents mean ± s.d. of n=10 Cy5-HA cryogels. Data in **d** represents mean ± s.d. of n=3 Cy5-HA cryogels and was compared using student’s t-test. Data in **f** represents mean ± s.d. of n=5 Cy5-HA cryogels. Data in **h** represents mean ± s.d. of n=3 HA cryogels and was compared using student’s t-test.

**Fig. S2.**
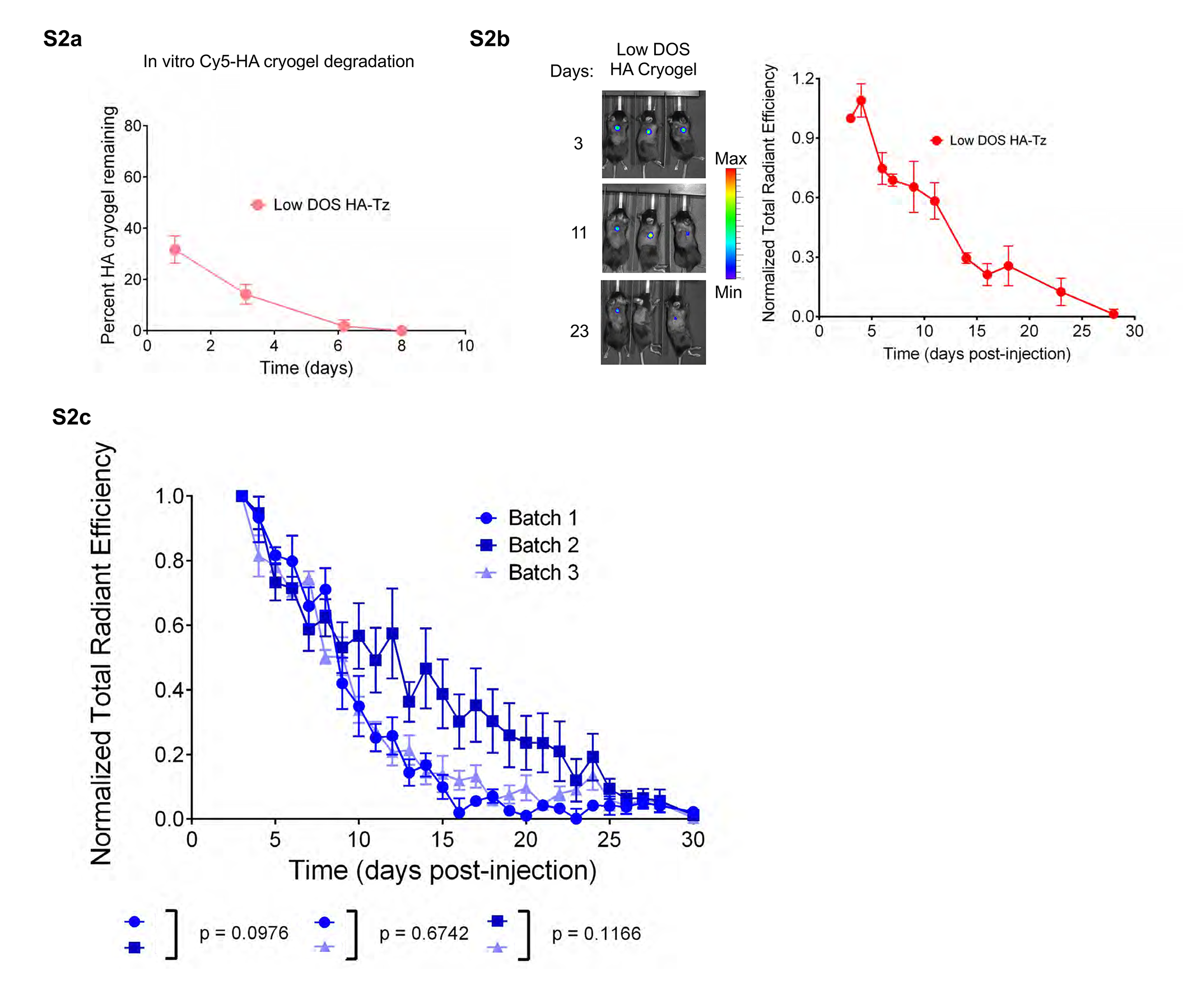
Supplementary Cy5-HA cryogel degradation characterization data (**a**) Measuring low-DOS Cy5-HA cryogel degradation in vitro by quantification of Cy5-signal in supernatant at pre-determined timepoints normalized to total Cy5-signal in supernatant across all timepoints. (**b**) Representative IVIS fluorescence images of gel degradation in mice and measuring low-DOS Cy5-HA cryogel degradation in vivo by quantification of total radiant efficiency normalized to the initial day 3 timepoint. (**c**) Measuring Cy5-HA cryogel degradation by quantification of total radiant efficiency normalized to initial day 3 timepoint. Data in **a** represents mean ± s.d. of n=4 HA cryogels. Data in **b** represents mean ± s.e.m. of n=4 HA cryogels. Data in **c** represents mean ± s.d. of n=4-5 HA cryogels and were compared using two-way ANOVA with Bonferroni’s multiple comparison test.

**Fig. S3.**
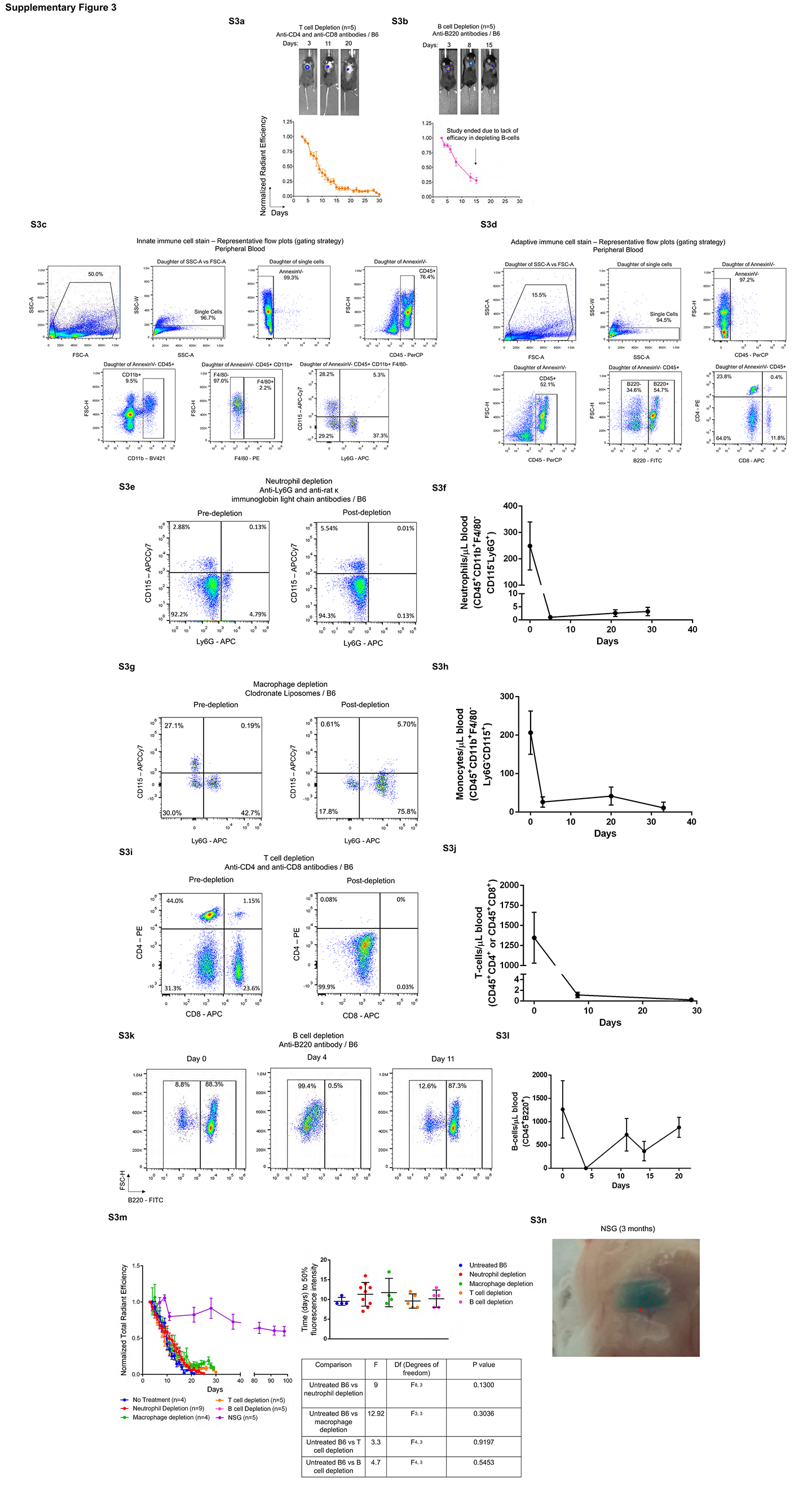
Supplementary Cy5-HA cryogel degradation in immunodeficient mice characterization data. Representative IVIS fluorescence images of gel degradation and quantification by measuring total radiant efficiency normalized to initial day 3 timepoint of (**a**) T cell depleted B6 mice and (**b**) B cell depleted B6 mice. **c-d** Representative gating strategy to determine identity of (**c**) innate immune cells and (**d**) adaptive immune cells in peripheral blood. (**e**) Representative flow cytometry plot of peripheral blood neutrophils pre- and post-administration of neutrophil depleted mice and (**f**) peripheral blood neutrophil concentration. (**g**) Representative flow cytometry plot of peripheral blood monocytes pre- and post-administration clodronate liposomes to mice and (**h**) peripheral blood monocyte concentration. (**i**) Representative flow cytometry plot of peripheral blood T cells blood pre- and post-administration of anti-CD4 and anti-CD8 antibody treatment to mice and (**j**) peripheral blood T cell concentration. (**k**) Representative flow cytometry plot of peripheral blood B cells blood pre- and post-administration of anti-B220 antibody treatment to B6 mice and (**l**) peripheral blood B cell concentration. (**m**) Overlay of normalized total radiant efficiency curves and time to 50% fluorescence intensity of untreated B6, neutrophil depleted, macrophage depleted, T cell depleted, and B cell depleted mice. (**n**) Photograph of a Cy5-HA cryogel retrieved from NSG mice 3 months post-injection. Data in **a**, **b** represents mean ± s.e.m. of n=5 and are representative of at least two separate experiments. Data in **f**, **h**, **j**, **l** represents mean ± s.d. of n=4-5. Data in **m** represents mean ± s.d. of n=4-9 and were compared using student’s t-test.

**Fig. S4.**
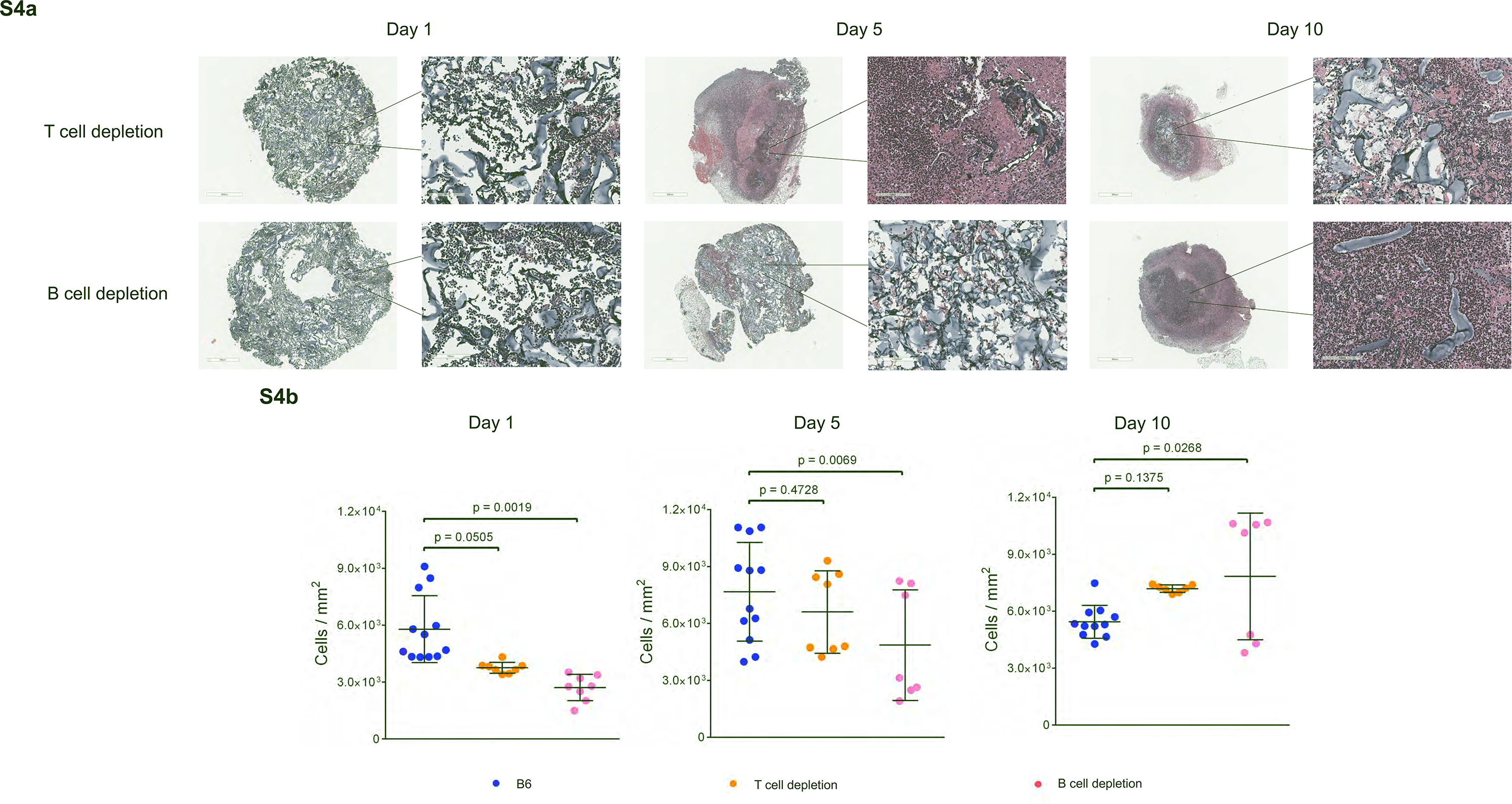
Supplementary histomorphometric analysis of Cy5-HA cryogels retrieved from T- and B-cell depleted mice (a) Hematoxylin and eosin (H&E) stain of explanted Cy5-HA cryogels from T cell depleted and B cell depleted mice at days 1, 5, and 10. Scale bar left = 800μm, scale bar right = 100μm. (b) Analysis of H&E stains to quantify cellular density in Cy5-HA cryogel. Data in b represents mean ± s.d. of n = 7-12 histological sections and was compared using student’s t-test.

**Fig. S5.**
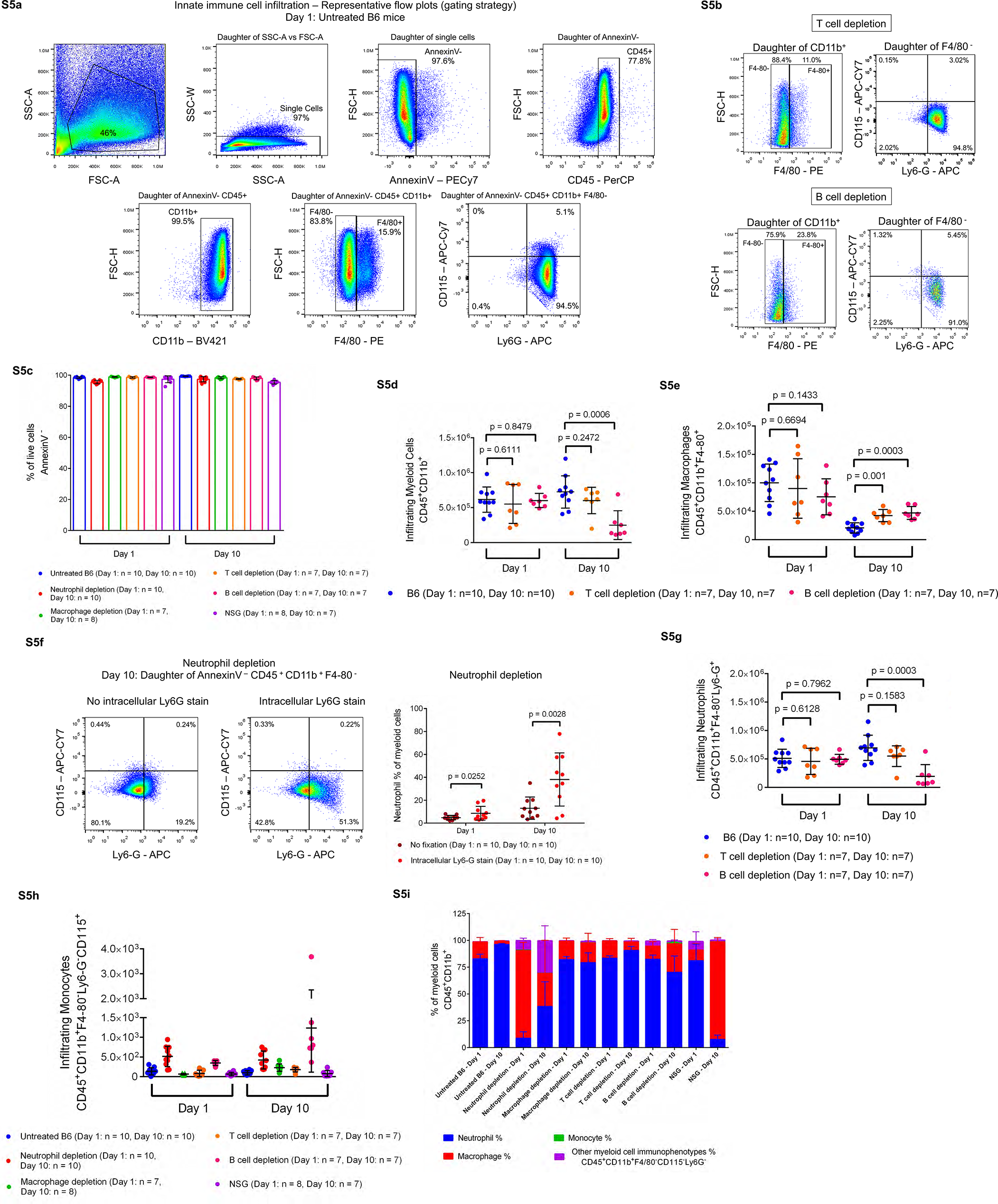
Supplementary analysis of myeloid cell infiltration of Cy5-HA cryogels retrieved from immunodeficient mice (a) Representative gating strategy to determine identity of innate immune cell infiltrates of HA cryogel. (b) Representative flow cytometry plots gated to determine cellular identity of CD45+ CD11b+ F4/80+ (macrophage) cells, CD45+ CD11b+ F4/80-Ly6G+ (neutrophil) cells, and CD45+ CD11b+ F4/80-Ly6G-CD115+ (monocyte) cells T cell depleted and B cell depleted mice. (c) Percent of AnnexinV-(live) cells within Cy5-HA cryogels one and ten days after implant from flow cytometry analysis. d-e Quantification of total number of (d) myeloid cells and (e) macrophages infiltrating Cy5-HA cryogels in untreated B6 mice, T cell depleted mice, and B cell depleted mice. (f) Representative flow cytometry plots from neutrophil depleted mice with and without intracellular Ly6G staining. Plotted data assessing neutrophils as a percentage of total myeloid cells (CD45+CD11b+) with and without intracellular Ly6G staining. (g) Quantification of total number of neutrophils infiltrating Cy5-HA cryogels in untreated B6 mice, T cell depleted mice, and B cell depleted mice. h,i Quantification of total number of (h) monocytes and (i) infiltrating immune cell lineages plotted as a percentage of myeloid cells in untreated, neutrophil depleted, macrophage depleted, T cell depleted, B cell depleted, and NSG mice. Data in e represents mean ± s.d. of n = 10. Data in c, d, e, g, h, i represents mean ± s.d. of n = 7-10 and are representative of at least two separate experiments. Data in d, e, f, g were compared using student’s t-test.

**Fig. S6.**
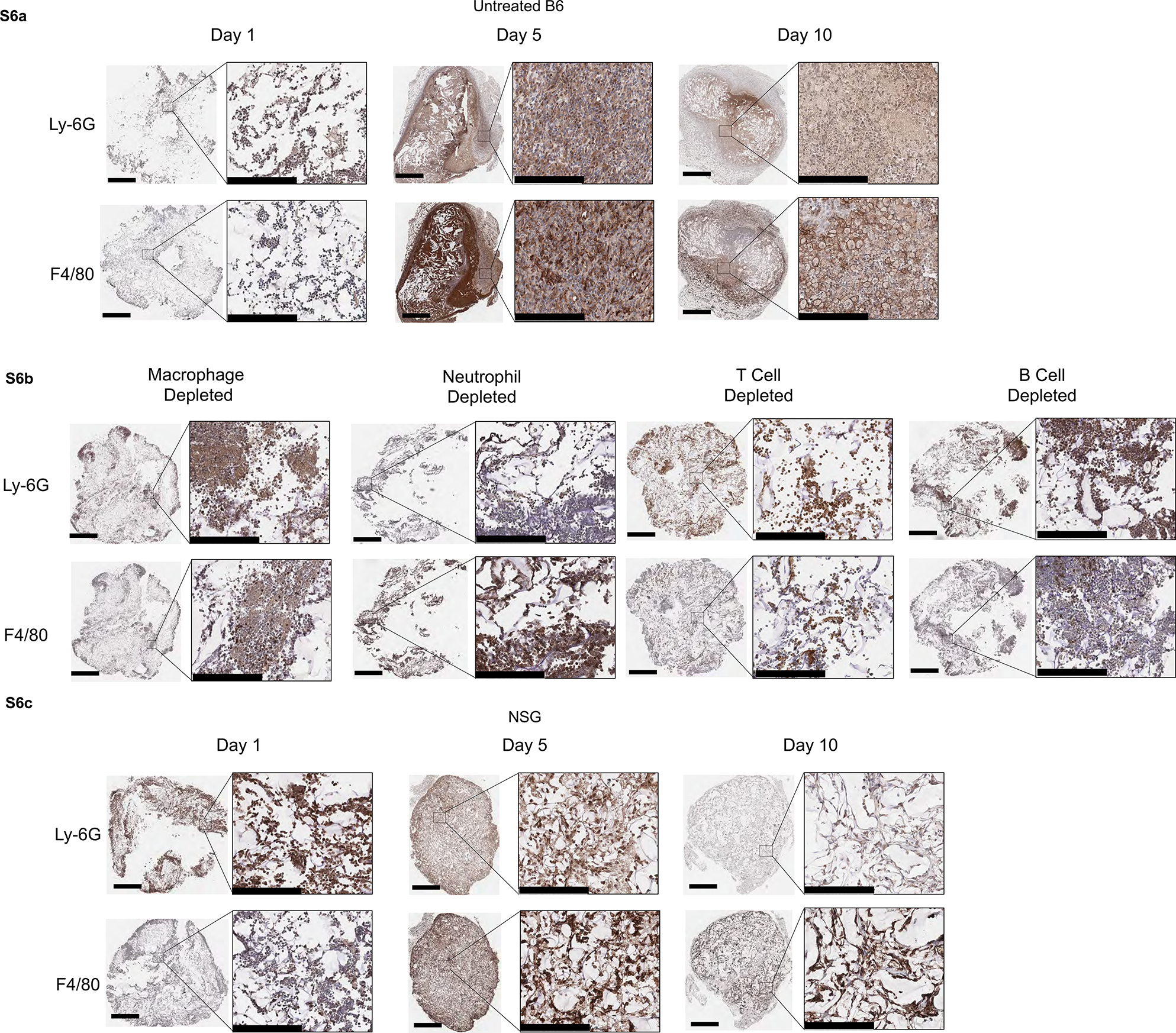
Supplementary Immunohistochemical staining of Cy5-HA cryogels retrieved from untreated B6 and NSG mice (a) Immunohistochemistry (IHC) staining for Ly6G (neutrophils, top, scale bar = 1mm) and F4/80 (macrophages, bottom, scale bar = 60μm) of Cy5-HA cryogels excised from untreated B6 mice 1-, 5-, and 10-days after injection. IHC was conducted on the same Cy5-HA cryogels as in Fig. 2e. (b) IHC staining for Ly6G (top, scale bar = 1mm) and F4/80 (bottom, scale bar = 60μm) of Cy5-HA cryogels excised from macrophage depleted, neutrophil depleted, T cell depleted, and B cell depleted B6 mice 1-day after injection. IHC was conducted on the same Cy5-HA cryogels as in Fig. 2e. (c) IHC staining for Ly6G (top, scale bar = 1mm) and F4/80 (bottom, scale bar = 60μm) of Cy5-HA cryogels excised from NSG mice 1-, 5-, and 10-days after injection. IHC was conducted on the same Cy5-HA cryogels as in Fig. 2e.

**Fig. S7.**
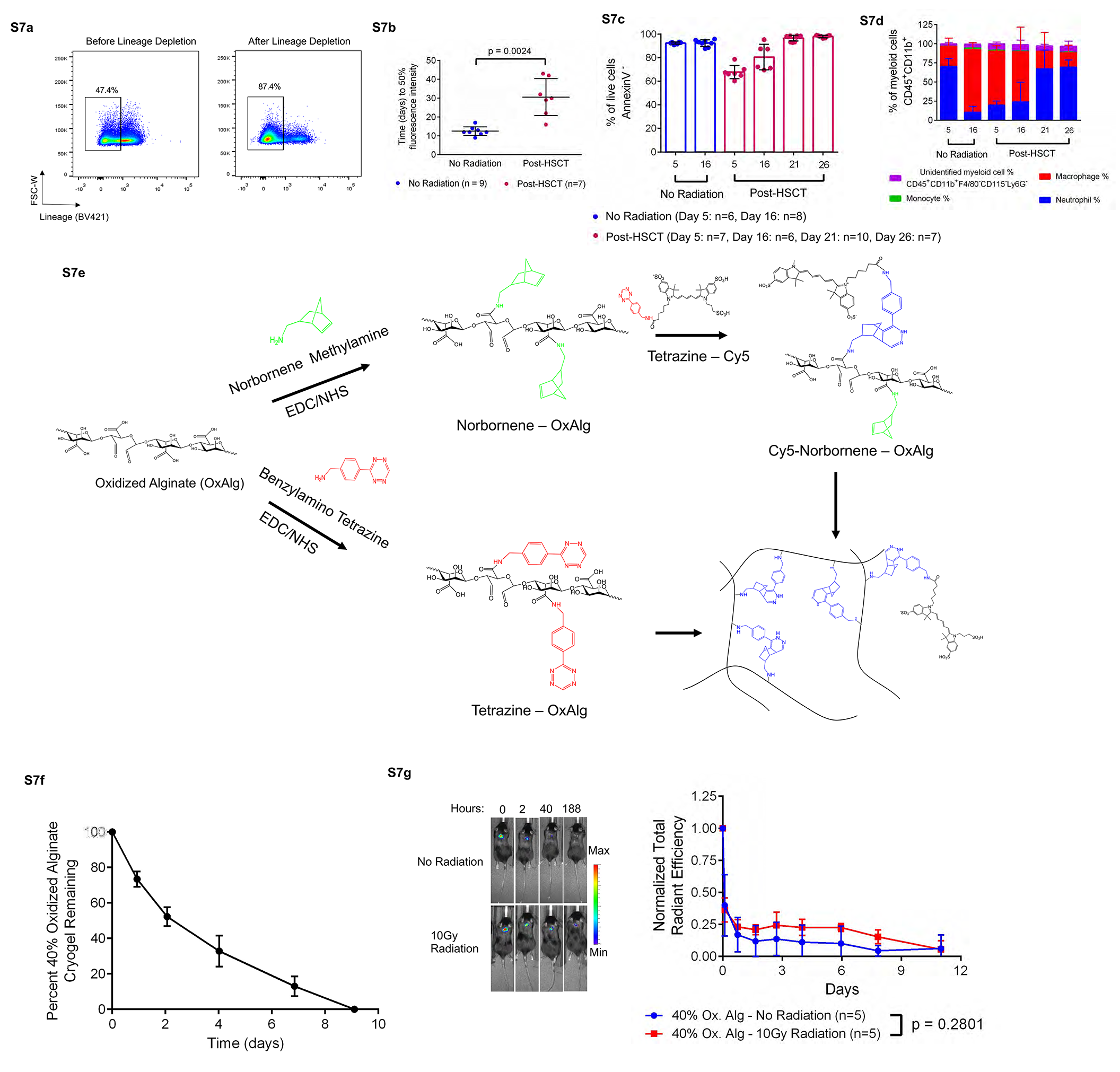
Supplementary data for degradation kinetics of HA cryogels post-HSCT Representative flow cytometry plots of bone marrow before and after lineage depletion. (**b**) Time to 50% fluorescence intensity of Cy5-HA cryogels in non-irradiated and post-HSCT mice. (**c**) Percent of AnnexinV-(live) cells within Cy5-HA cryogels 5- and 16-days post injection in non-irradiated mice and 5-, 16-, 21-, and 26-days post-injection in post-HSCT mice. (**d**) Infiltrating immune cell lineages plotted as a percentage of myeloid cells in non-irradiated and post-HSCT mice. (**e**) Schematic for tetrazine (Tz) and norbornene (Nb) functionalization of oxidized alginate (OxAlg), Cy5 functionalization of Nb functionalized OxAlg, and crosslinking of Tz functionalized HA with Cy5 functionalized OxAlg. (**f**) Measuring Cy5-OxAlg cryogel degradation in vitro by quantifying the Cy5-signal in supernatant at pre-determined timepoints normalized to total Cy5-signal in supernatant across all timepoints. (**g**) Representative in vivo imaging system (IVIS) fluorescence images of gel degradation in mice and measuring Cy5-tagged 40% oxidized alginate cryogel degradation in vivo by quantification of total radiant efficiency normalized to initial 2-hour timepoint. IVIS Images are on the same scale and analyzed using Living Image Software. Data in **b** represents n=7-9 Cy5-HA cryogels, is representative of at least two separate experiments and were compared using student’s t-test. Data in **c, d** represents mean ± s.d. of n=6-10 Cy5-HA cryogels and is representative of at least two separate experiments. Data in **f** represents mean ± s.d. of n=5 Cy5-OxAlg cryogels. Data in **g** represents mean ± s.e.m. of n=5 Cy5-OxAlg cryogels and were compared using two-way ANOVA with Bonferroni’s multiple comparison test.

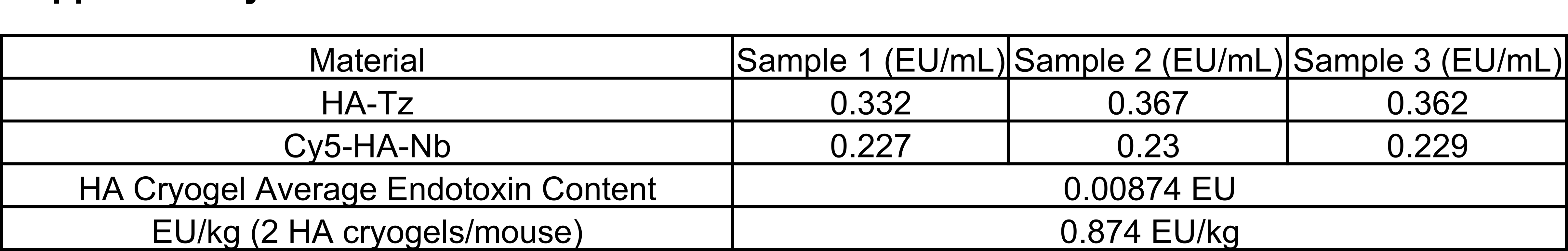

**Supplementary Table 1**
| Material | Sample 1 (EU/mL) | Sample 2 (EU/mL) | Sample 3 (EU/mL) |
| --- | --- | --- | --- |
| HA-Tz | 0.332 | 0.367 | 0.362 |
| Cy5-HA-Nb | 0.227 | 0.23 | 0.229 |
| HA Cryogel Average Endotoxin Content | 0.00874 EU |  |  |
| EU/kg (2 HA cryogels/mouse) | 0.874 EU/kg |  |  |

**Supplementary Table 2**
| Depletion Type | Depletion Agent | Dose, Administration Route, Frequency | Depletion Efficiency |
| --- | --- | --- | --- |
| Neutrophil Depletion | Anti-mouse Ly6G antibody (1A8, Bio X Cell)<br>Anti-rat kappa immunoglobulin light chain antibody (MAR 18.5, Bio X Cell) | 25µL anti-mouse Ly6G - IP administration - Everyday for 1 week<br>50µL anti-mouse Ly6G - IP administration - Everyday after first week<br>50µL anti-rat kappa immunoglobulin light chain - IP administration - Every other day | Consistent - 98% Depletion of neutrophils in peripheral blood<br>(Supplemental 5b,5c) |
| Macrophage/Monocyte Depletion | Clodronate Liposomes (Liposoma) | 100µL - IP administration - 2x/week | Consistent - 80-95% Depletion of monocytes in peripheral blood<br>(Supplemental 5d, 5e) |
| T-cell Depletion | Anti-mouse CD4 antibody (GK1.5, Bio X Cell)<br>Anti-mouse CD8α antibody (2.43, Bio X Cell) | 400µg anti-mouse CD4 - IP administration - 2x/week<br>400µg anti-mouse CD8α - IP administration - 2x/week | Consistent - 99% Depletion of T-cells in peripheral blood<br>(Supplemental 5f, 5g) |
| B-cell Depletion | Anti-mouse B220/CD45R antibody (RA3.31, Bio X Cell) | 400µg anti-mouse B220/CD45R - IP administration - 2x/week | Transient - 99% Depletion of B-cells in peripheral blood 4 days after start of depletion. Full reconstitution of B-cells by 3 weeks.<br>(Supplemental 5h, 5i) |

